# PDX models reflect the proteome landscape of pediatric acute lymphoblastic leukemia but divert in select pathways

**DOI:** 10.1101/2019.12.18.880401

**Authors:** Anuli C. Uzozie, Enes K. Ergin, Nina Rolf, Janice Tsui, Amanda Lorentzian, Samuel S. Weng, Lorenz Nierves, Theodore G. Smith, C. James Lim, Christopher A. Maxwell, Gregor S.D. Reid, Philipp F. Lange

**Affiliations:** Department of Pathology, University of British Columbia, Vancouver, BC, Canada; Michael Cuccione Childhood Cancer Research Program, BC Children’s Hospital Research Institute, Vancouver, BC, Canada; Department of Pediatrics, University of British Columbia, Vancouver, BC, Canada; Department of Molecular Oncology, British Columbia Cancer Research Centre, Vancouver, BC, Canada

**Keywords:** leukemia, N terminome, phosphorylation, proteome, xenograft

## Abstract

Murine xenografts of pediatric leukemia are known to accurately recapitulate genomic aberrations. How this translates to the functional capacity of the proteome is unknown. Here, we studied global protein abundance, phosphorylation, and proteolytic processing in 11 pediatric B- and T-cell acute lymphoblastic leukemia patients and 19 corresponding xenografts. Protein level differences that stratified pediatric disease subtypes at diagnostic and relapse stages were largely recapitulated in xenograft models. Patient xenografts lacked multiple human leukocyte antigens, and complement proteins, and presented incomplete response mechanisms to the host immune system which is absent in the murine model. The dominant expression of MKI67 and cell cycle proteins indicated a high proliferative capacity of xenografted cells residing in the spleen. Structural genomic changes and mutations found in patients were reflected at the protein level. The post-translational modification landscape is shaped by leukemia type and host and only to a limited degree by the patient of origin. This study portrays how genomic and host factors shape protein and post-translational modification landscapes differently, and confirms murine patient-derived xenograft as competent model system while highlighting important areas of diverging biology.

## Introduction

Established clinical phenotypes and genomic alterations characterize pediatric acute lymphoblastic leukemia (ALL), the most common hematologic malignancy in early childhood. The diverse prognosis associated with ALL includes morphological bone marrow changes and a transformed immunophenotype. Pediatric ALL is molecularly different from adult ALL, and presents specific primary alterations and oncogenic drivers^1,2^. In early B-cell ALL, this includes genetic rearrangements and copy number alterations in *ETV6*-*RUNX1* and *BCR*-*ABL*, MLL rearrangements, hypodiploid and hyperdiploid karyotypes^3^. Mutational events in *STIL*-*TAL1*, *CDKN2A/CDKN2B*, *NOTCH1*, and *PTEN* are characteristic to T-cell acute lymphoblastic leukemias. These core genetic alterations, combined with other cooperating oncogenic drivers contribute to leukemogenesis.

The impact of animal models on leukemia research cannot be overstated^4^. Non-obese diabetic (NOD) severe combined immunodeficient mice (SCID) mice with deletion in interleukin 2 receptor gamma chain (IL2Rgamma), lack functional B- and T-cells, have diminished natural killer cells, lack a functional complement system and have been proven to excellently engraft primary patient leukemia cells^4,5^. Although highly adequate for studying clinical drug assessments and blast-stromal interactions^4^, patient derived xenograft (PDX) models using immunodeficient mice do not enable investigation of leukemia immune cell interactions. Xenografts from B-cell ALL, T-ALL, and acute myeloid leukemia (AML) maintain patient-specific leukemogenic profile pertaining to the transcriptome, MRD marker expression, immunophenotype, and chromosomal aberrations^6–10^. DNA methylation profiling likewise revealed strong similarities in the epigenome of matched primary and xenograft blasts^11^.

The major determinants of disease phenotypes and most common drug targets are proteins, which unlike genes and mRNAs, directly modulate cell function and response to perturbation. Following translation, proteins can be diversely modified, and made capable to simultaneously regulate multiple processes and pathways in separate cellular locations^12^. Formation of functionally different proteoforms^13^ by post-translational modification (PTM) is frequently in diseases, including childhood cancers^14^. Protein phosphorylation and irreversible proteolytic processing are among the main PTMs that regulate cellular processes and contribute to the development and progression of cancer. The entirety of protein functional molecules and associated mechanisms broadly impacted in leukemic subtypes is yet to be understood. Such knowledge is crucial to determine if disease stratification based on proteome features can be attained, how well this complements known genomic subtypes, and what biological mechanisms provide stable molecules for precise therapeutic targeting. Functional and drug sensitivity studies in PDX models of leukemia can only be extrapolated to patients if the relevant pathways, protein networks and modifications are conserved between patient and murine model. Cases of conflicting protein abundance have been reported in human and xenograft leukemia prototypes. The use of PDX models for studying selected somatic mutations such as *IDH1/2* or *TET2* is considered risky due to divergent manifestation of these defects in patient and PDX models^8^. Also, conventional ALL xenograft models were found unsuitable for clinical studies targeting JAK-STAT signaling pathway, due to their non-expression of thymic stromal lymphopoietin (TSLP) protein^15^. The efficacy of JAK inhibitors is highly dependent of the activation of cytokine receptor-like factor 2 (CRFL2) by TSLP.

Mass spectrometry-based proteomic techniques facilitate rapid, extensive, reproducible, and accurate characterization of cellular proteomes ^13,16^. Global protein abundance, protein phosphorylation and proteolytic processing can be comprehensively studied using enrichment methods coupled to mass spectrometry-based proteomic techniques. Protein phosphorylation is vital for cell signaling, regulating a multitude of cellular processes^17^. Kinase inhibitors are employed as targeted treatment for cancers, including pediatric malignancies^18–20^. These drugs target known kinase families and have proven effective against specific disease subtypes with chromosomal rearrangements that dysregulate kinase signaling. For example, Imatinib, a tyrosine kinase inhibitor, targets the BCR-ABL fusion protein, and is a front-line regimen for patients with pediatric Philadelphia chromosome-positive (Ph+) ALL^18^. Global assessment of phosphorylation requires relatively large specimen to achieve good coverage of phosphorylation events. This constitutes a major challenge for phospho-proteomic techniques, limiting their application to clinical biopsies. However considerable methodological improvements now enable phosphoproteomic analysis of sub-mg quantities at moderate depth^21–23^. Limited proteolytic processing is an irreversible enzymatic modification that regulates physiological and pathological processes through inactivation or changes in function, location, and stability of proteins. It is mediated by proteases/peptidases, and results in new stable protein fragments (proteoforms) with a distinct protein N terminus^24^. We recently developed a new enrichment procedure (High-efficiency Undecanal-based N Termini Enrichment, HUNTER) to extensively profile the N terminome of minimal disease samples^24,25^. In addition to profiling proteolytic proteoforms, we and others have shown that the N terminome provides a reliable repertoire to verify the existence of truncated proteins, including stable fragments that are established biomarkers^26^ and identify their functional relevance in biological processes such as apoptosis^25,27–29^.

A proteomic investigation of pediatric ALL prior and after transplantation in immunodeficient mouse models will reveal proteome signatures characteristic of leukemic subtypes, and benchmark how effectively PDX models replicate the primary leukemic proteome. ALL-related PTMs would need to be conserved in xenograft models to correctly replicate relevant protein structure, functions, and interactions. Our results provide the first proteomic landscape of matched patient and PDX pediatric ALL, that substantiates the genomic profile of patient and xenograft ALL subtypes. Using a diverse clinical cohort we show that leukemic bone marrow blasts have a remodeled proteome distinct from mononuclear cells dominant in patients lacking leukemic blasts. In comparison to leukemia cell lines, patient-derived xenografts more closely replicated the protein profile of patients. Xenografts further retained protein-level presentation of gene mutations and structural rearrangements evident in patients. We confirmed protein and PTM differences in cellular pathways and processes that are lacking or compromised between patient and PDX models.

## Results

### Proteome differentiates pediatric B-ALL and T-ALL at diagnosis and relapse stages from non-leukemic cells

Leukemic cells from 13 samples comprising 8 pediatric B-ALL patients (4 diagnosis, 4 relapse, 1 relapse progression), 3 pediatric T-ALL patients (2 diagnostic, 1 relapse progression), and 1 pediatric patient with T-cell lymphoblastic lymphoma (T-LBL) (Table 1) were each transplanted in NOD/SCID/IL2 gamma-receptor null (NSG) mice. Thirteen primary leukemia samples, 19 corresponding xenograft leukemia, 2 non-leukemic samples, and 4 pediatric leukemia cell lines (B-ALL: 697, 380; T-ALL: DND41, PEER), were analyzed using multiple mass spectrometry-based proteomics strategies. To control for the effect of blast count in patient bone marrow cell population, two samples originating from patients with less than 10% leukemic blasts infiltration to the bone marrow compartment (higher proportion of normal hematopoietic mononuclear cells) were included in the study (Fig. 1A and Table 1). The scheme depicted in Fig. 1B illustrates the study workflow used to investigate total protein abundance, proteolytic processing and phosphorylation events in pediatric human and xenograft leukemia subtypes. The number of blood and bone marrow mononuclear cells attainable from pediatric patients is limited. For this study we had access to 0.6 – 5 million primary cells per patient. Minimal starting protein amounts mandated the use of highly sensitive workflows with minimum sample loss. Starting from 60 µg protein in crude cell lysate, we identified 6396 proteins (peptide and protein FDR = 0.01) (Fig. 1C), and 3531 phosphosites (localization probability > 0.75) in 3378 peptides (Fig. 1C). Likewise, 3853 N termini (FDR = 0.01) were quantified from 30 µg starting protein amounts, with more representation from free (experimentally dimethylated, 58%) compared to acetylated (42%) N Termini (Fig. 1C). We assessed the precision of our quantification by determining the coefficient of variation (CV) between repeat injections. Protein-level measurements for patients, and PDX had average CVs below 12% (Supplementary Fig. 1A). N Termini were quantified with average peptide level CVs of 23% in patients and 25% in PDX respectively (Supplementary Fig. 1B). Our methods therefore support robust and sensitive investigation of the proteome of childhood acute lymphoblastic leukemia with acceptable quantitative precision.

**Table 1:**
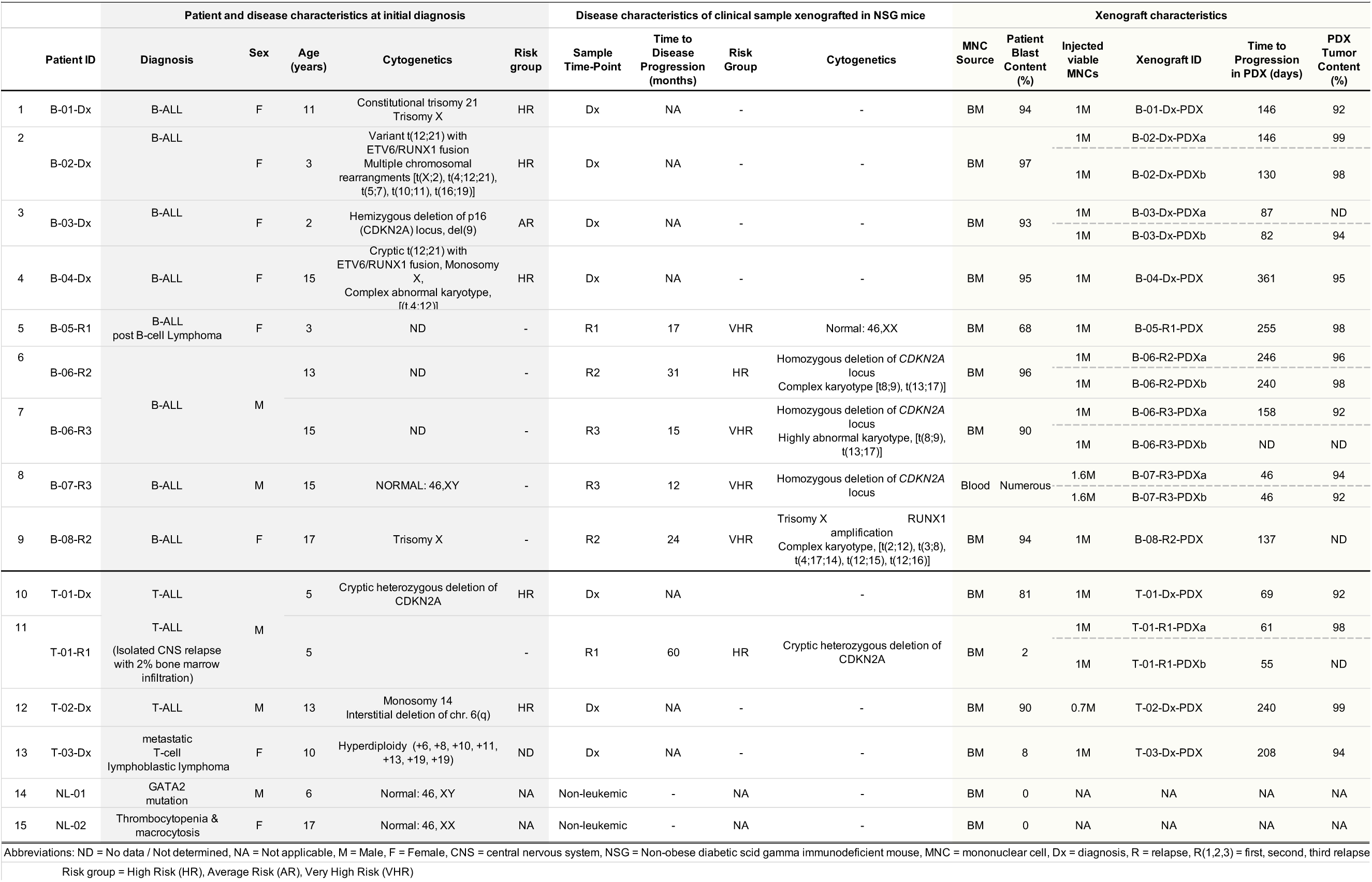
Clinical information on Patients and corresponding NSG Xenografts

**Figure 1.**
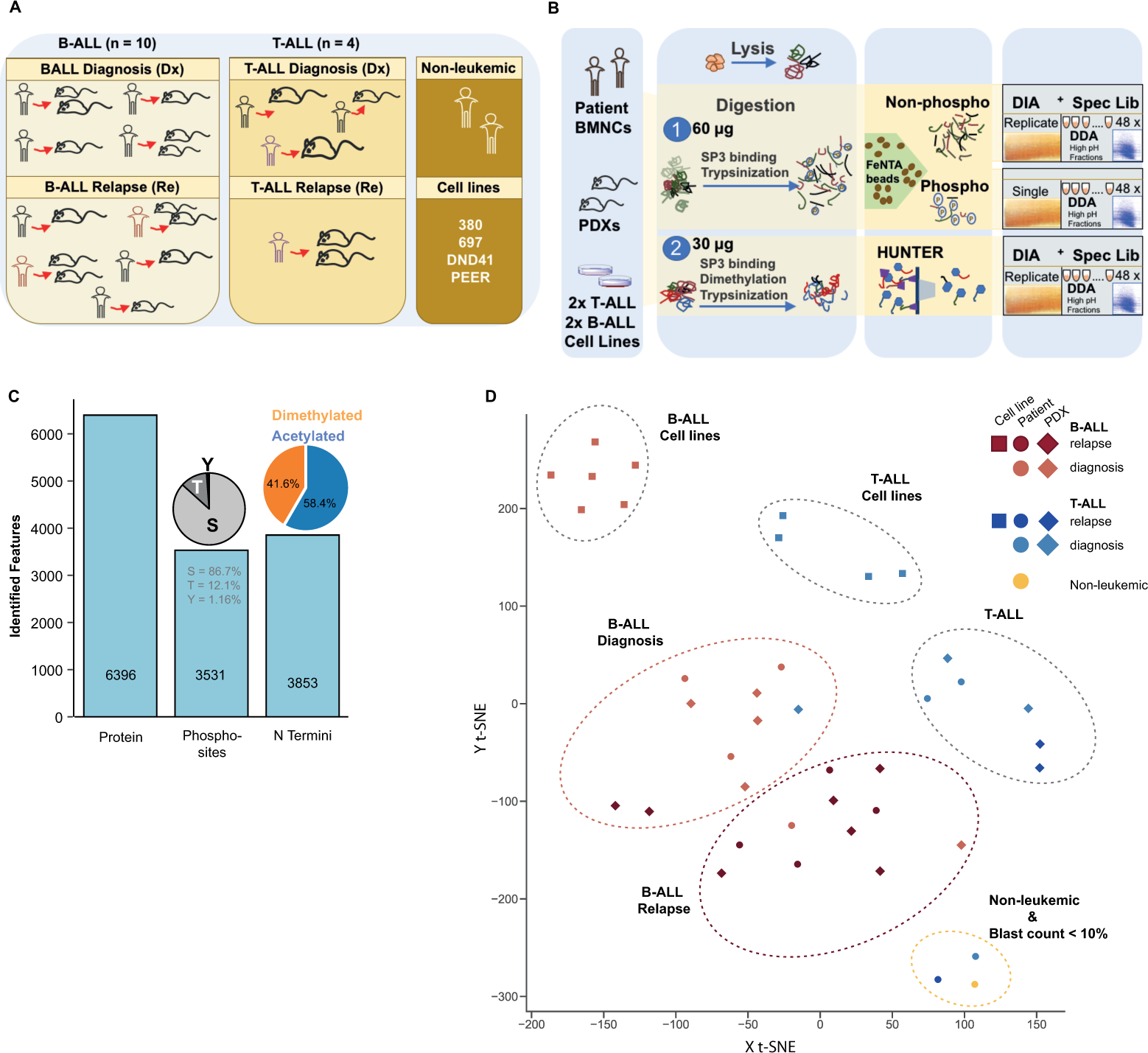
Proteomics stratifies pediatric ALL subtypes. A. Composition of samples including number of childhood leukemia subtypes and disease stages, non-leukemic group, and leukemic cell lines analyzed. Same colours are maintained for ALL patient samples collected at different disease stages. B. Detailed workflow for investigation of total protein, N Terminome, and phosphoproteome of pediatric ALL from minimal protein starting amount (60 µg protein for total and phosphoproteome study, 30 µg for N termini study). Proteome features were measured using Data Independent Acquisition (DIA) mass spectrometry methods, and analyzed with spectral libraries generated using combined information from DIA analysis and Data Dependent Analysis (DDA) of high pH fractionated sample pools. C. Summary of identifications at total protein (6,396), phosphosite (3,531), and N termini levels (3,853) respectively. D. T-distributed Stochastic Neighbor Embedding, t-SNE, plot following unsupervised analyses on average protein intensities (N = 5,554) and K-means clustering on reduced dimensions from t-SNE. Clusters depict protein-level similarities and differences between model organisms (13 patients, 19 PDXs, 4 cell lines), disease subtypes (8 B-ALL, 3 T-cell leukemia), and disease stages (7 diagnosis, 6 relapse).

Increased bone marrow cellularity and blast cell population correlated with a loss of mature myeloid, erythroid, and lymphoid cell populations (Table 2). The bone marrow morphology of normal and diseased patients differed as expected. To investigate if the proteome of non-malignant samples could represent characteristics of a non-leukemic pediatric bone marrow phenotype devoid of blast cells, we examined the representation of biological pathways and processes in the non-leukemic and diseased categories - B-ALL patients, B-ALL xenografts, T-ALL patients, T-cell leukemia patients with low blast cell population, and T-cell leukemia xenografts. The overlap of proteins quantified in each group is shown in Supplementary Fig. 1C for B-cell leukemia and in Supplementary Fig. 1D for T-cell leukemia. Using Metascape^30^, enrichment analysis was performed comparing proteins quantified in each of the listed categories. This revealed that the spectrum of processes and pathways enriched in the “non-leukemic” and “low blast count” samples were different from those enhanced in patient and xenograft leukemia (Supplementary Figs. 2 and 3, Supplementary Table 1 and 2). The proteome landscape of pediatric ALL patients we studied clearly captures pathological features of bone marrow leukemic blasts before and after xenotransplantation.

**Table 2:**
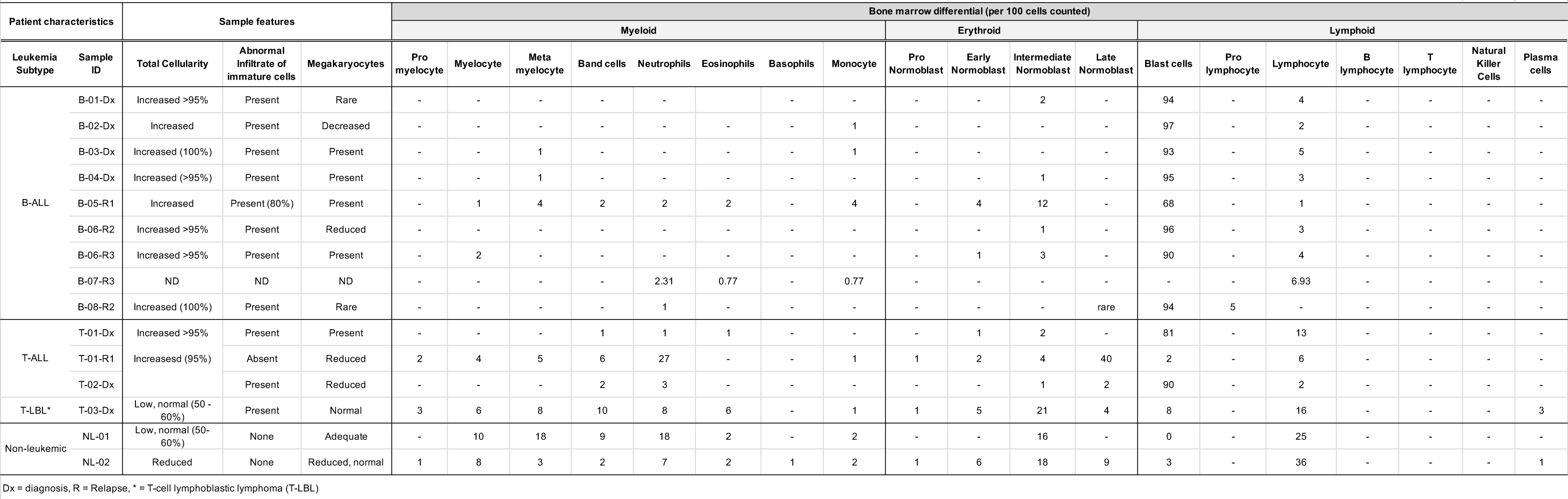
Bone marrow morphology for clinical cohort

**Figure 2.**
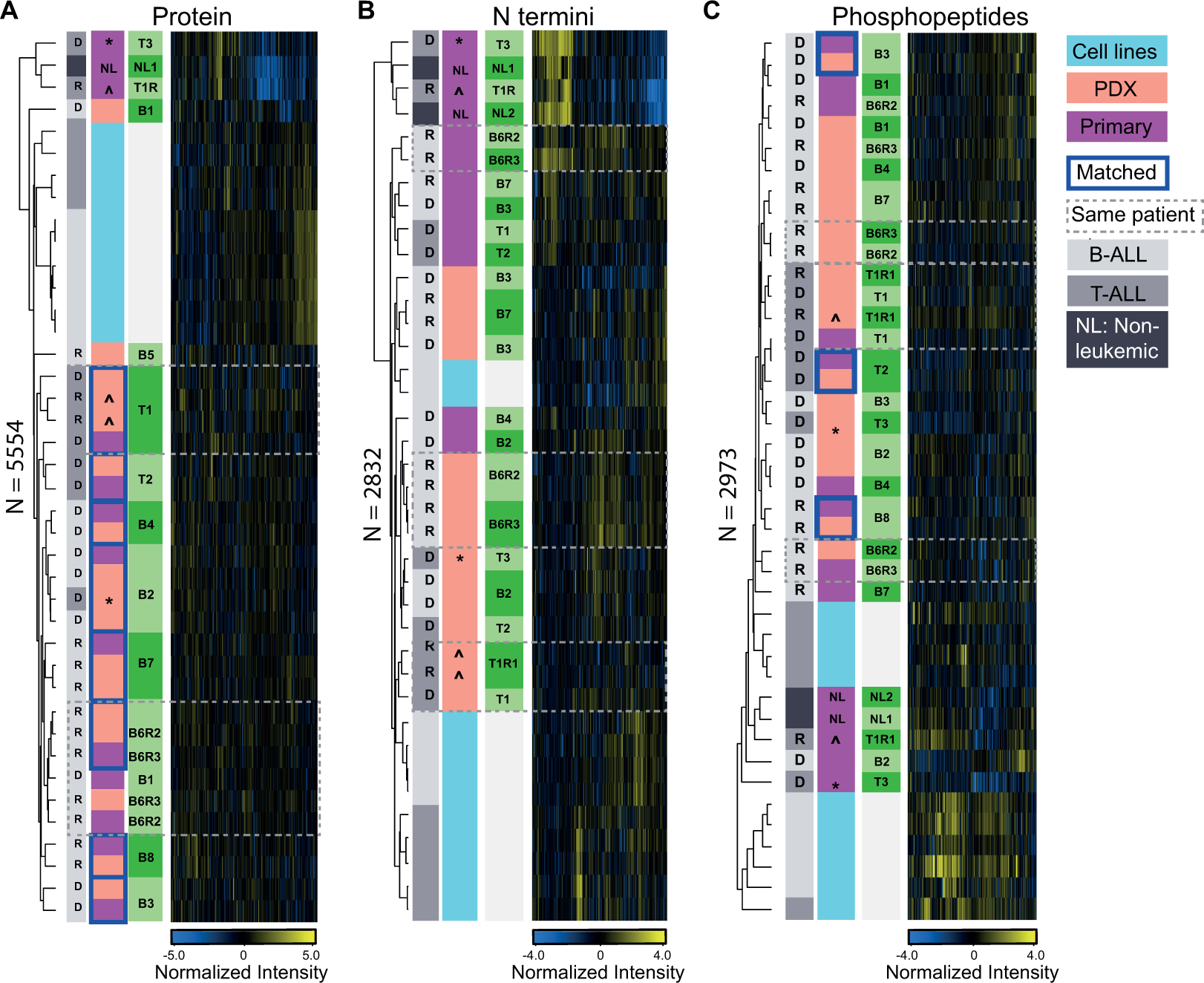
Pediatric ALL proteome landscape in patients, PDXs and leukemic cell lines. A, B, C. Hierarchical clustering map following unsupervised analyses on (A) 5554 quantified proteins, (B) 2832 N termini, and (C) 2973 phosphopeptides. Values plotted in heat maps are z-score normalized. Paired patient and xenograft samples are highlighted with blue border lines. The symbols ^ and * represent two samples, T-01-R1 and T-03-Dx, with low disease involvement of the bone marrow (<10% of blasts), and their corresponding PDXs.

**Figure 3.**
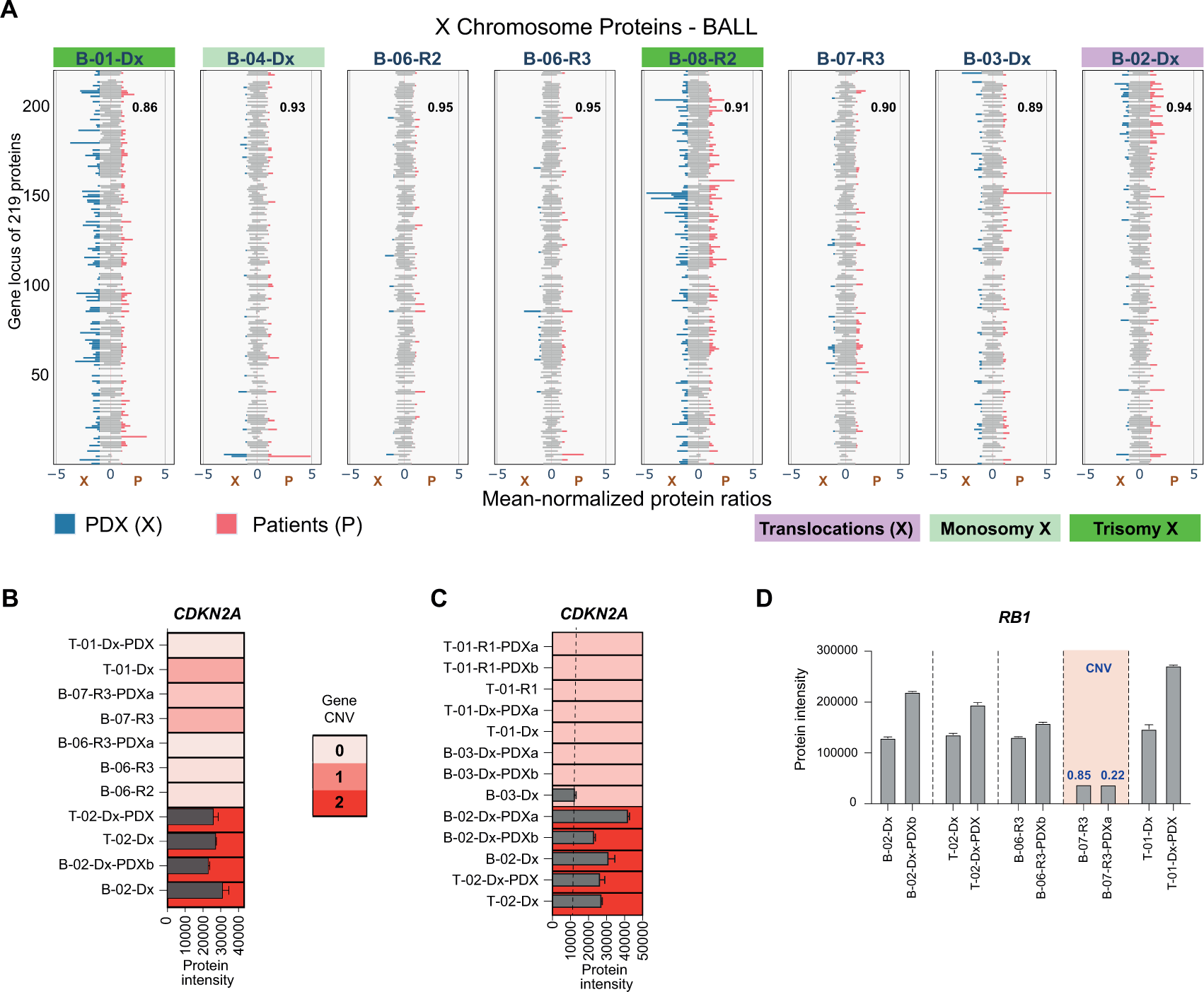
Pediatric ALL proteome reflects consequences of genomic changes. A. Mean-normalized protein intensities of 219 quantified protein products from genes on chromosome X in 8 patients (P, pink bars) and in their matched PDXs (X, blue bars). Average intensities are reported for multiple PDXs engrafted with the same patient leukemia. Patients with structural defects in their X chromosome are highlighted – translocation in purple, monosomy X in light green, and trisomy X in dark green. Pearson correlation coefficient score for each matched patient and PDX pair is provided inset. B. *CDKN2A* copy number determined by targeted gene sequencing in matched patient and PDXs, and CDKN2A protein level in same patient and xenograft samples. C. CDKN2A protein level in all samples with one or two copies of *CDKN2A*, as confirmed by clinical cytogenetics and/or targeted gene sequencing. D. *RB1* copy number variation determined by targeted gene sequencing, and RB1 protein level in same paired patient and xenograft samples. Bar plots are mean protein abundance from two DIA technical replicates, and error bars represent the standard deviation from the mean. CNV = copy number variation, and is highlighted in blue.

### Proteomes of xenograft ALL most closely resemble the matched primary patient specimen

Unsupervised T-distributed Stochastic Neighbor Embedding (t-SNE) dimensionality reduction followed by K-means clustering on protein abundance measurements categorized samples in distinct groups (Fig. 1D). Notable cluster groups were evident for non-leukemic samples, B-ALL samples at diagnosis and relapse, T-ALL cases, and leukemic cell lines respectively. Strikingly, samples from the two T-cell leukemia patients with very low blast infiltration of the bone marrow (blast count < 10%), consistently associated more with the non-malignant sample. (Fig. 1D and Fig. 2A). PDX largely co-clustered with their matched primary patient specimen after dimensionality reduction (Fig. 1D) and pairwise hierarchical clustering (Fig. 2A). Mice that received blasts from patients with limited blast count (<10%, T-01-R1 and T-03-Dx respectively) developed overt leukemia, and the protein profile of the expanded blasts associated more closely with leukemic samples than with their primary samples. The average correlation scores (Spearman rank correlation) for protein abundance in T-ALL patient and corresponding xenograft mice was 0.83 (SD = 0.06) (Supplementary Fig. 4A), with lower correlations (average = 0.67, SD = 0.06)) observed between primary leukemias with lower disease burden (blast count <10%) and matched PDXs. B-cell leukemia samples had on average 0.88 (SD = 0.02) Spearman rank correlation score (Supplementary Fig. 4B). Proteins involved in known signaling pathways affected in pediatric leukemia, including JAK-STAT and PI3K pathways, showed a similar or stronger correlation between primary and PDX leukemias (Supplementary Figs. 4A and 4B). Pediatric PDX models principally recapitulate the protein abundance landscape of patients.

**Figure 4.**
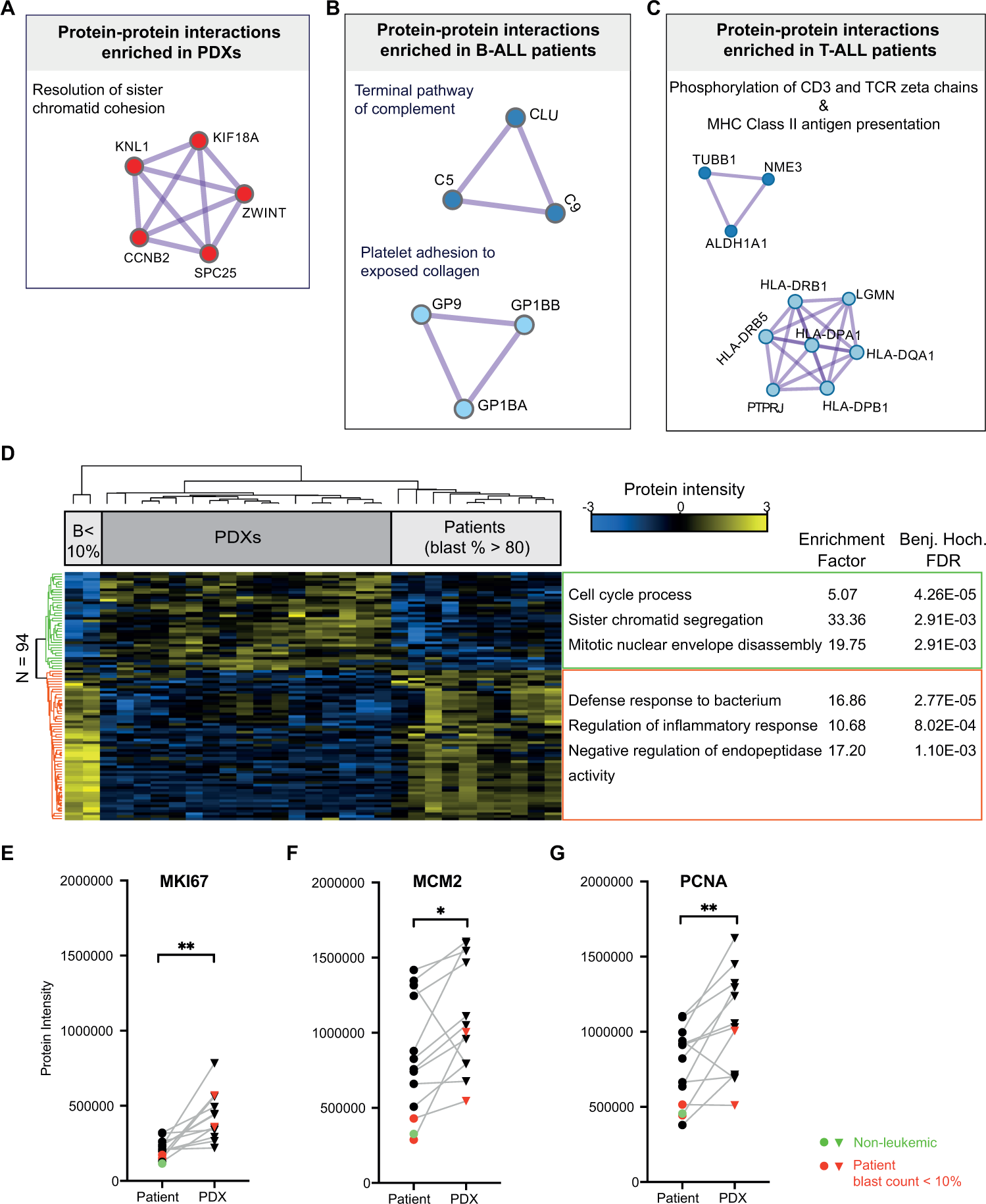
PDXs demonstrate differences in cell proliferation and immune response processes compared to patients. A, B, C. Proteins unique to either PDX or host, and found to be involved in established interactions and functional roles in (A) both B-ALL and T-ALL xenografts, (B) B-ALL patients, and (C) T-ALL patients (See methods for description of analysis in Metascape). D. Supervised hierarchical clustering of differentially regulated proteins (Two-tailed Student’s t-test, FDR < 0.05) in all patients compared to all PDXs. Heatmap depicts z-score normalized intensities for 94 proteins. Processes and pathways enriched in proteins in each of two distinct clusters are displayed (Fisher exact test, Benjamini-Hochberg FDR < 0.02). Enrichment test was performed using annotations for 5554 total proteins as background. Patient samples with reduced bone marrow disease involvement (blast count, B < 10%) are indicated. E-G. Two-tailed paired Student’s t-test on (E) MKI67, (F) MCM2 and (G) PCNA protein abundance in 12 matched patient and PDX pairs. Samples from the non-leukemic patient and from the two patients with blast infiltration less than 10% (2) are highlighted in green and red respectively. Symbols * and ** denote p-values of 0.001 (MKI67), 0.03 (MCM2), and 0.006 (PCNA).

Unlike protein level clustering, unsupervised hierarchical clustering of protein N termini abundance grouped samples primarily by host and leukemia type (Fig. 2B). Although matched patient and PDX did not co-cluster, a positive correlation (B-ALL average = 0.62, SD = 0.06 and T-ALL average = 0.60, SD = 0.03) was retained between individual patients and their corresponding xenografts (Supplementary Fig. 5). Of note, the N terminome of the two patients with low leukemic burden had little or no correlation with their PDX recipients (average Spearman correlation = 0.13, SD = 0.04). Compared to the proteome landscape, the N Terminome landscape of pediatric ALL is not as well replicated in immunocompromised mice and appears primarily shaped by host factors.

**Figure 5.**
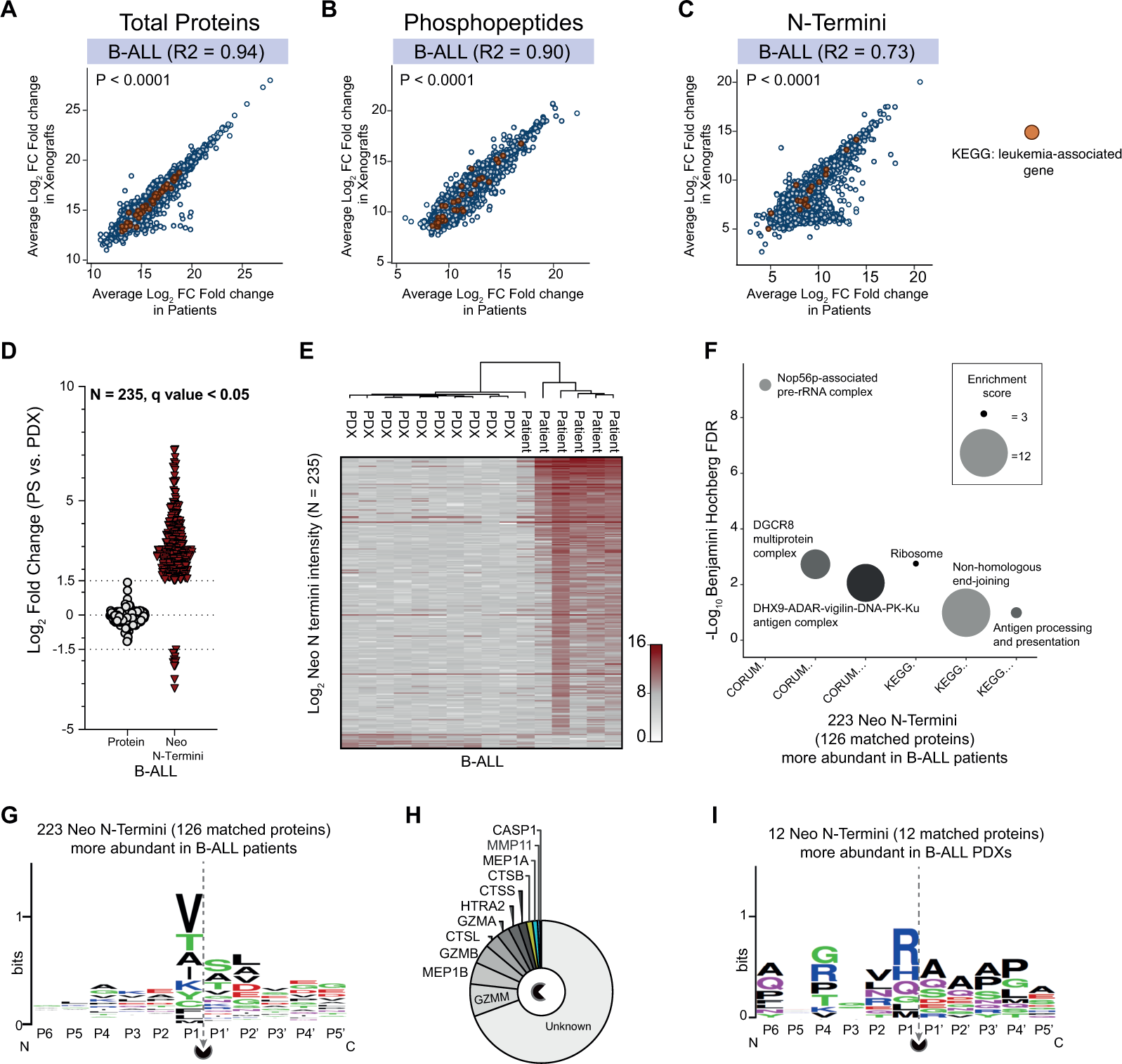
Neo N-terminal PTMs show differences in B-ALL patients and PDXs that are absent at protein level. A, B, C. Correlation on average Log_2_ fold changes between patients and PDXs for (A) proteins (N = 5554), (B) phosphopeptides (N = 2973) and (C) N termini. Proteins annotated to KEGG leukemia-associated genes are coloured in dark brown filled circles for proteins (N = 54), phosphopeptides (N = 35) and N Termini (N = 17). D. Proteins (N = 235) that remained stable between B-ALL patients and PDXs (Log_2_ fold change < ±1.5) but differed in neo N termini abundance in xenografts and patients (Log_2_ fold change >±1.5, unpaired two-tailed Student’s t-test q value < 0.05). E. Log_2_ abundance plot of 235 differentially regulated neo N termini in patients and corresponding PDXs. F. Enriched pathway (KEGG) and protein complexes (CORUM) mapped to protein-matched upregulated neo N termini (N = 126 proteins from 223 neo N termini). Circle size represents the enrichment score of the category term. G. Sequence pattern detected for 223 neo N termini enriched in B-ALL patients. H. Curated proteases with cleavage sites associated with the sequence patterns identified in 223 neo N termini. I. Sequence pattern detected for 12 neo N termini enriched in B-ALL xenografts.

From minimal clinical material (60 µg protein lysate), we quantified 3513 class I phosphosites (localization probability > 0.75) from 3,378 phosphorylated sequences, corresponding to 1,450 phosphoproteins. Unsupervised hierarchical clustering showed no close association between matched primary and PDX leukemias (Fig. 2C).

Compared to ALL expanded cells and non-leukemic bone marrow cells, the proteomes of patient-derived xenograft ALL most closely resemble that of their patient at protein level, while at PTM level model- and disease-specific similarities exceed paired patient-PDX relationships.

### Xenografts recapitulate proteome response to structural genomic changes in patients

Quantified proteins were matched to gene chromosome location in UniProt^31^. For each chromosome, in a given disease subgroup, within a defined model, we calculated the mean protein abundance in samples. Protein intensities were then normalized to the respective mean protein value. Normalized protein ratios were aligned based on gene location and visualized. Multiple patients with distinct protein patterns for chromosome X (Fig. 3A) had structural alterations in chromosome X (Table 1). In particular, patients B-01-Dx and B-08-R2, had trisomy X, patient B-04-Dx had a monosomy X, while patient B-02-Dx had a translocation between chromosome X and 2, t(X;2). Despite a loss of one X chromosome, patient B-04-Dx retained protein features similar to all other patients lacking alterations in chromosome X, irrespective of biological gender category. The correlation scores (Fig. 3A, inset) confirmed these protein changes were stably reflected in matched patient xenografts.

We next validated whether specific genomic aberrations in patients were maintained in their corresponding xenograft leukemia. Targeted next-generation sequencing (NGS) was performed on a subset of six paired primary and PDX samples using the recently published Oncomine Childhood Cancer Research Assay^32^. Mutations detected in B-ALL and T-ALL leukemias include single nucleotide alterations, gene fusions, copy number variations, and are detailed in Table 3. An *ETV-RUNX1* fusion in patient B-02-Dx at diagnosis was also detected in the equivalent xenograft leukemia (B-02-Dx-PDXb). While loss of one copy of *SOCS2* and *TET2* was apparent in only B-02-Dx-PDXb. Single nucleotide variations in ARID1A (Gln563Ter) and in KRAS (Lys117Arg) were consistently retained in patient B-05-R1 and in its paired xenograft leukemia. Additional variations including loss of one copy each of *CDKN2A* and *CDKN2B* occurred in B-05-R1-PDX. For three cases, B-06, B-07, and T-01, the exact alterations were preserved in their respective xenograft leukemias. These results support previously known findings that genetic alterations in primary leukemia are mostly retained in PDX models, however, clonal expansion in xenograft leukemia increases the chances for detecting additional tumor suppressor or leukemia driver mutations^7,11^.

**Table 3.** Genomic alterations detected in patient and corresponding xenograt leukemia by next generation sequencing using the OCCRA gene panel (Lorentzian et al., 2019)

| Disease Type | Sample Origin | OCCRA panel NGS |
| --- | --- | --- |
| **1**<br>B-ALL Dx | **Primary**<br>B-02-Dx | *ETV6-RUNX1* fusion<br>SNV: SUZ12 lys690fs, 0.03 |
|  | **PDX**<br>B-02-Dx-PDXb | *ETV6-RUNX1* fusion<br>CNV: SOCS2 (0.83), TET2 (0.97) |
| **2**<br>B-ALL R1 | **Primary**<br>B-05-R1 | SNV: ARID1A (Gln563Ter, 0.33), KRAS (Lys117Arg, 0.35) |
|  | **PDX**<br>B-05-R1-PDX | SNV: ARID1A Gln563Ter (0.49), KRAS Lys117Arg (0.52)<br>CNV: FASLG (3.67), CDKN2A (0.85), CDKN2B (0.96) |
| **3**<br>B-ALL R3 | **Primary**<br>B-06-R3 | SNV: JAK1 (Val658Phe, 0.48)<br>CNV: CDKN2A (0.07), CDKN2B (0.07), PTCH1 (0.79), EBF1 (1.0) |
|  | **PDX**<br>B-06-R3-PDXb | SNV: JAK1 (Val658Phe, 0.99)<br>CNV: CDKN2A (0), CDKN2B (0), PTCH1 (0.76), EBF1 (1.0) |
| **4**<br>B-ALL R3 | **Primary**<br>B-07-R3 | SNV: KRAS Gln61His (0.13)<br>CNV: CDKN2A (0.52), RB1 (0.85), P53 (1.37) |
|  | **PDX**<br>B-07-R3-PDXa | SNV: KRAS Gln61His (0.4)<br>CNV: CDKN2A (0.33), RB1 (0.22), P53 (1.1) |
| **5**<br>T-ALL Dx | **Primary**<br>T-02-Dx | *STIL-TAL1* fusion<br>SNV: PTEN (Leu247fs, 0.2) |
|  | **PDX**<br>T-02-Dx-PDX | *STIL-TAL1* fusion |
| **6**<br>T-ALL Dx | **Primary**<br>T-01-Dx | CNV: CDKN2A (0.6) |
|  | **PDX**<br>T-01-Dx-PDX | CNV: CDKN2A (0.03) |

We then determined if our proteomics data could associate protein patterns to CNVs and SNVs validated by targeted NGS. Cyclin-dependent-kinase inhibitor 2A (CDKN2A) is a well-known tumor suppressor protein that is inactivated in pediatric leukemia^6,33^. Deletion in *CDKN2A* on chromosomal region 9 of p21 was detected at a rate of 62% in our sequenced B- and T-ALL sample cohort (12 samples) (Table 3). Patients with a complete loss of *CDKN2A* (CNV = 0) verified in our sequenced cohort lacked detectable protein amounts (Fig. 3B). Interestingly, patients with one-copy loss of *CDKN2A* had no protein or reduced protein levels in contrast to patients with both copies of the gene (Fig. 3C).

For patient B-02 with deletions in one copy of *SOCS2* and *TET2* respectively (Table 3), no substantial depletion in levels of SOCS2 and TET2 was detected at peptide and protein FDR of 0.01. The same patient had a cryptic *ETV6-RUNX1* (t(12:21)) fusion. Both ETV6 and RUNX1 protein levels did not differ significantly in B-02 in comparison with other diagnostic B-ALL samples.

Reduced protein levels following a hemizygous copy loss in the retinoblastoma protein, RB1, were conserved between patient B-07 and its corresponding PDX, analyzed by targeted NGS. Compared to all sequenced sample pairs, RB1 protein abundance was significantly decreased in B-07 patient and xenograft (Table 4 and Fig. 3C). Additionally, RB1 protein levels were similar in the multiple xenograft samples from patients B-02, B-06, and B-07, which were not analyzed by targeted NGS (Supplementary Fig. 6).

**Figure 6.**
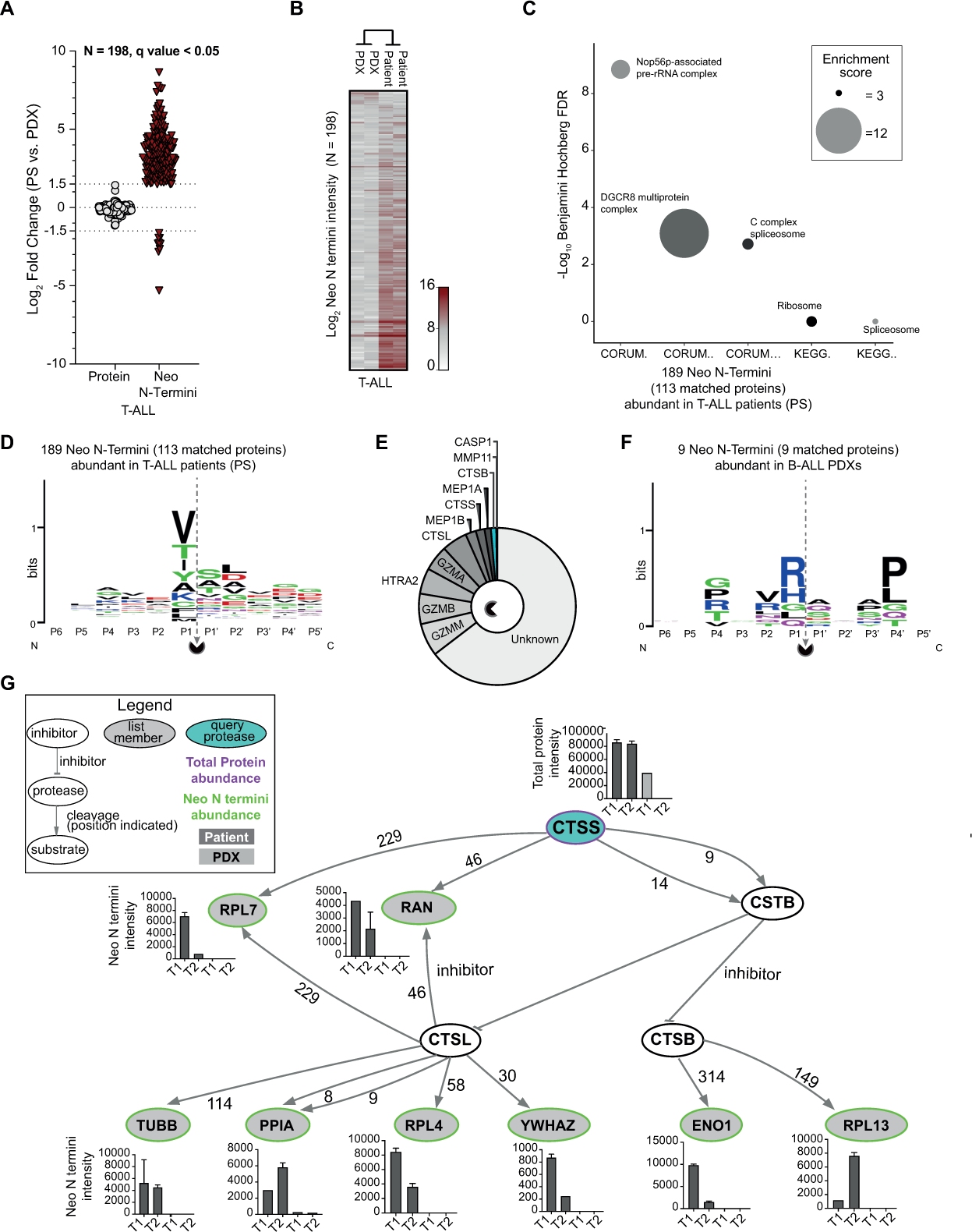
Neo N-terminal PTMs show differences in T-ALL patients and PDXs that are absent at protein level. A. Proteins (N = 198) that remained stable between T-ALL patients and PDXs (Log_2_ Fold change < ±1.5) but differed in neo N termini abundance in PDX and host (Log_2_ fold change >±1.5, unpaired two-tailed Student’s t-test q value < 0.05). B. Log_2_ abundance plot of 166 differentially regulated neo N termini in patients and corresponding PDXs. C. Enriched pathway (KEGG) and protein complexes (CORUM) mapped to protein-matched neo N termini (N = 113 proteins from 189 neo N termini). Circle size represents the enrichment score of the category term. D, E. Sequence pattern detected for 189 neo N termini enriched in B-ALL patients and (E) curated proteases with cleavage sites associated with the identified patterns. F. Sequence motif for 9 neo N termini enriched in B-ALL PDXs. G. Protease web plot of CTSS interaction network. CTSS (query protease, purple border) mean protein abundance, and Neo N termini mean abundance for list members (proteins with neo N termini in query list, green border) (N =198) are plotted in bar graphs. Error bars indicate the standard deviation from the mean for replicate DIA-mass spectrometry measurements. The amino acid following the protease N terminal cleavage site (amino acid P1’) is specified for each protease-substrate path.

### Proliferation and immune response processes differ between PDXs and patients

We next evaluated if, amongst the global similarity of patient and PDX proteome landscapes, individual processes differ. We first compared biological processes and pathways enriched in proteins that were uniquely identified in patients or in PDXs (Supplementary Figs. 7A and 7B). Multiple immune-related, metabolic and extracellular pathways were significantly enriched in both B- and T-ALL patients when compared with xenograft ALL (Supplementary Figs. 7C and 8A). In xenografts, processes linked to cell division and regulation of cyclin-dependent protein serine/threonine kinase activity were enhanced (Supplementary Fig. 8B). Protein-protein interaction enrichment analysis (See methods) further identified pathways containing protein complexes that might be disrupted or functionally compromised by altered expression of complex members in either patients or PDXs. For ALL xenografts, a set of 5 enriched interacting proteins were functionally assigned to the pathway “Resolution of Sister Chromatid Cohesion (R-HSA-2500257)”, a key step in mitotic anaphase during cell division (Fig. 4A). In B-ALL, components of two pathways were identified. Proteins CLU, C9, and C5, functionally linked to the Reactome pathway “Terminal pathway of complement” (R-HSA-166665). And, GP9 (CD42a), GP1BA (CD42b), and GP1BB(CD42c), with functional annotation to “Platelet adhesion to exposed collagen” (R-HSA-75892) (Fig 4B). While in T-ALL “Phosphorylation of CD3 and TCR zeta chains, R-HSA-202427 and MHC Class II antigen presentation, R-HSA-2132295” were the most enriched pathways represented by interacting proteins (Fig. 4C). The absence of these proteins could be linked to the lack of a human immune system in NSG mice.

**Figure 7.**
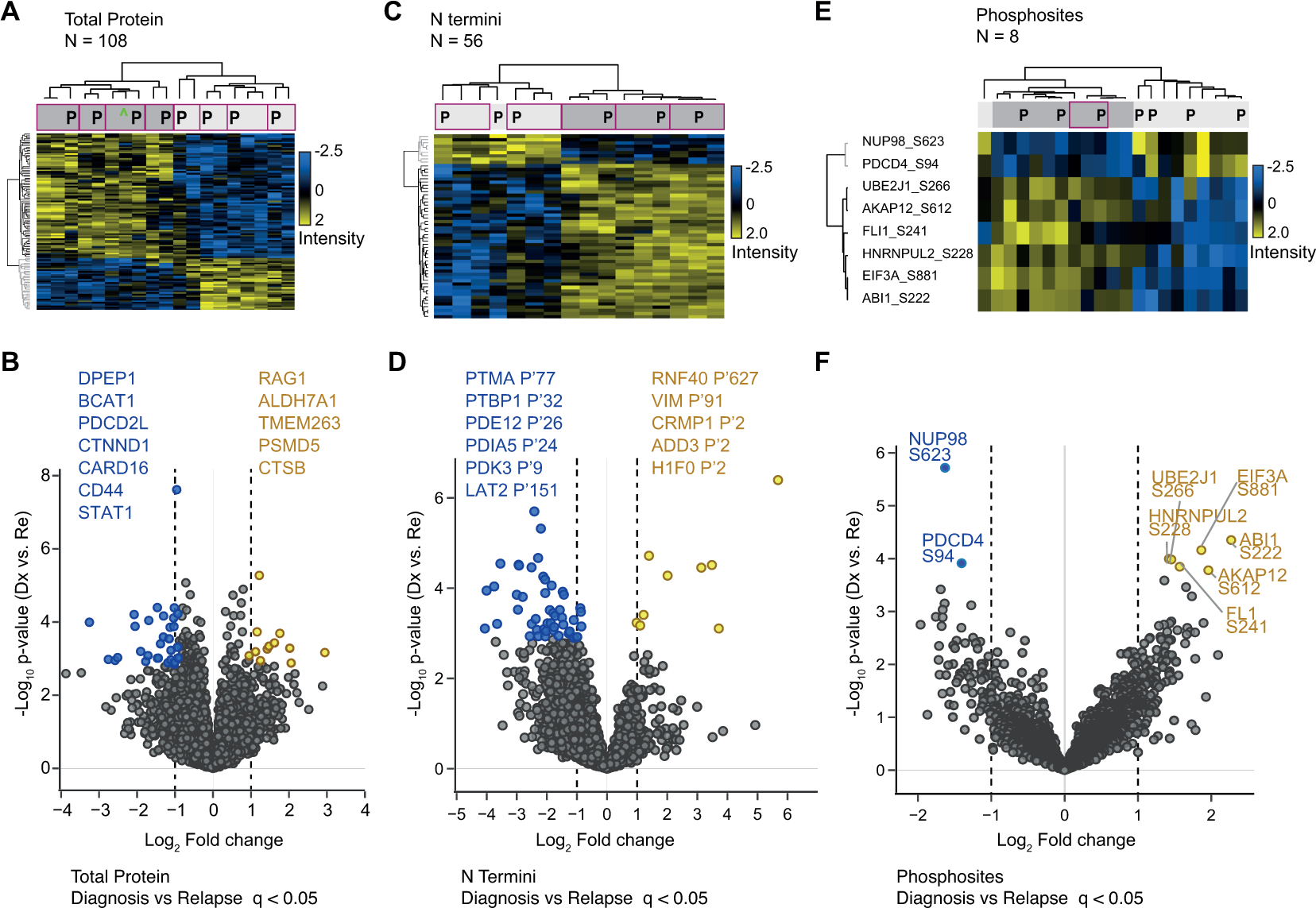
PDX reflect proteome changes that distinguish diagnosis and relapse conditions in B-ALL patients. A. Heatmap on normalized intensities of 108 proteins that significantly distinguished diagnosis from relapse B-ALL conditions (supervised hierarchical clustering, unpaired Student’s t-test q value < 0.05). B. Fold change and p-value profiles of the top differentiating proteins (q value < 0.05) in diagnosis and relapse samples. C. Supervised hierarchical clustering on 56 N termini that significantly distinguished (unpaired Student’s t-test q value < 0.05) diagnosis from relapse samples. D. N termini with significant differences between diagnosis and relapse time-points (q value < 0.05) are highlighted in a scatter plot. E. Supervised hierarchical clustering on 8 phosphosites with differential abundance in diagnosis and relapse cases (ANOVA test, q value < 0.05) F Scatterplot shows significantly dysregulated phoshosites in diagnosis and relapse patients (q value < 0.05). Purple border highlights paired patient (P) and PDX samples.

**Figure 8.**
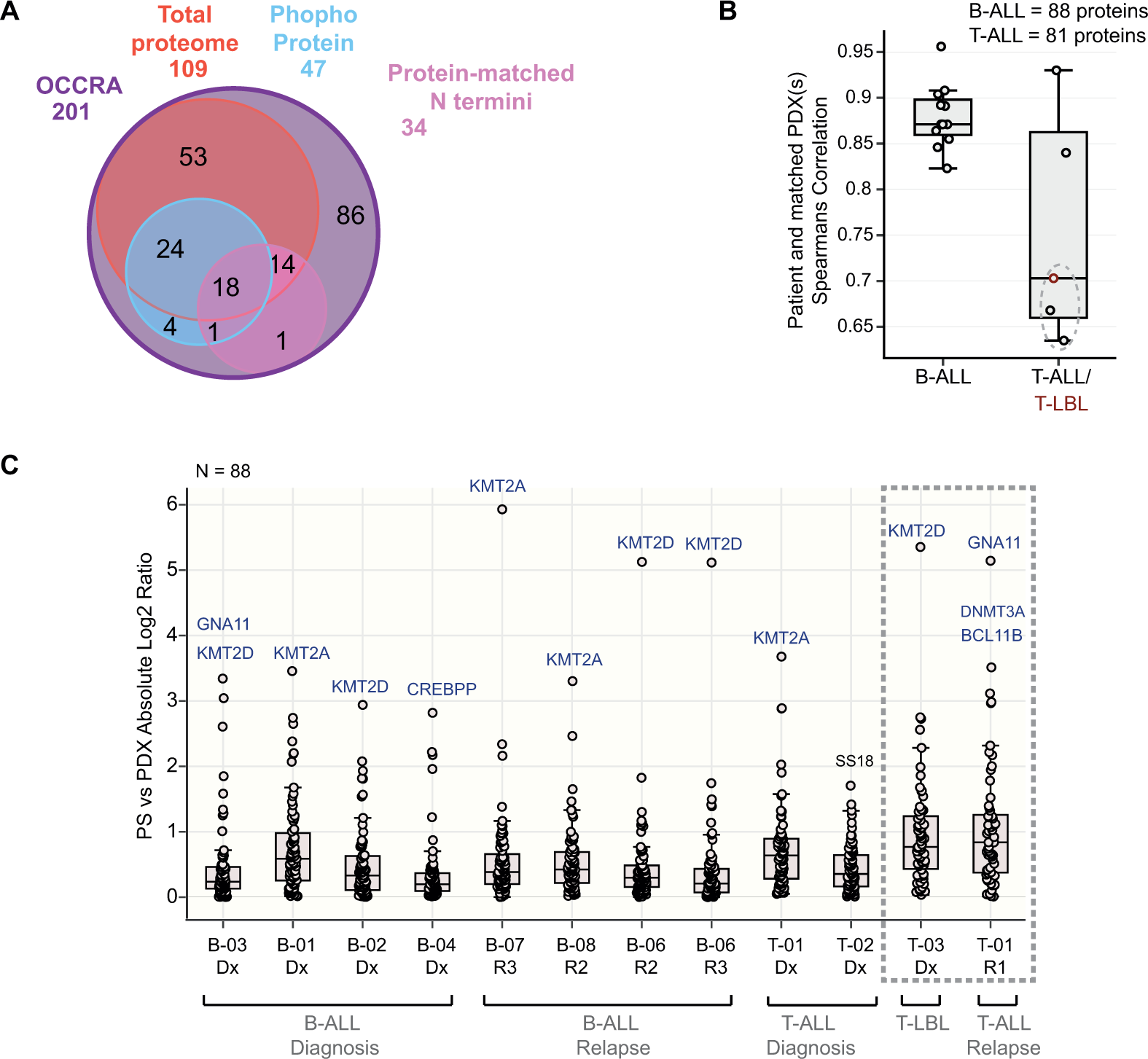
Pediatric ALL-related gene expression changes are reflected in PDX. A. Summary of 201 OCCRA panel genes profiled in total protein (109), phosphoprotein (47), and matched protein N termini (34) data from clinical samples. B. Spearman correlation scores for quantified OCCRA proteins in paired patient and PDX leukemia samples (B-ALL = 88 proteins, T-ALL = 81 proteins). Points highlighted with dashed semi-circles represent values from comparing protein abundance in patients with low blast count (<10%) and their matched PDXs: T-01-R1 and T-01-R1-PDXa, T-01-R1 and T-01-R1-PDXb, T-03-Dx and T-03-Dx-PDX respectively. C. Absolute Log_2_ ratios of protein intensities (N = 88) in 12 ALL patients and their corresponding PDX. Proteins with highest variation between both models are highlighted in blue. Values form patients with reduced bone marrow involvement (<10% blasts) and their corresponding PDXs are highlighted in dashed border lines.

We next identified processes enriched in proteins showing significantly altered protein abundance between patient and xenografted blasts. A two-sample Student’s t-test was employed to compare protein abundance between all primary ALL and all PDX leukemia samples. The 94 proteins with significantly altered abundance (FDR < 0.05) between patients and PDXs clustered in two main groups (Fig. 4D). A Fisher exact test to determine non-random Gene Ontology (GO) process associations between these proteins and all proteins quantified revealed that proteins with lower abundance in PDX specifically associated with immune and inflammatory cellular response. This included proteins S100A12, S100A9 and S100A8, with major roles in cytokine promotion and chemokine production. In contrast, proteins related to cell cycle and mitosis had significantly higher levels in PDXs when compared to patients. These results concurred with enriched patterns attributed to proteins detected or missing between PDXs and murine hosts.

To confirm if cell proliferation was indeed increased in xenografts when compared to patient leukemia, we quantified the abundance of known proliferation markers MKI67, PCNA, and MCM2, across all paired samples (Figs. 4E-G). MKI67, PCNA and MCM2 levels were significantly increased in all PDX compared to their primary counterparts (paired Student’s t-test, p < 0.05). PCNA and MCM2 levels had a downward trend in xenograft leukemia from sample B-01-Dx.

### Post-translational modifications are largely conserved but proteolysis shows select differences between patient and PDX not evident at the protein level

Limited proteolysis by proteases results in an ensemble of stable protein fragments with neo N Termini distinct from that of the unprocessed protein. Also, cellular signalling is highly dependent on kinase phosphorylation activities. Overall, we found that phosphorylation strongly correlates (R^2^ = 0.90, p-value < 0.0001) between the patient and xenograft groups while N termini abundance correlates moderately (R2 = 0.73, p-value < 0.0001) (Figs. 5A-C, Supplementary Figs. 9A-9C). We next investigated if proteins that are stable between patients and PDX show altered post-translational modification. For each ALL subtype, proteins that did not significantly change between patient and PDXs (Student’s t-test, q value > 0.05) were evaluated for significant differences (Student’s t-test, q value < 0.05, fold change >= ±3) in phosphorylation sites or in neo N termini. No significant changes in phosphosite and phosphopeptide levels that contrasted protein abundance were detected, while a subset of protein N termini indicated differences in proteolytic processing.

To evaluate the altered proteolytic processing in detail we then characterized protease-generated neo N termini (see Methods for definition) in ALL subtypes using TopFIND^34,35^ and TopFINDER^36^. Comparing patients and PDX of B-cell origin, we identified 235 changing neo termini in proteins that remained stable (Figs 5D and 5E). Specifically, 224 neo termini were higher in patients while 12 were elevated in PDX. To evaluate the functional relevance of these neo termini, biological pathway (KEGG) and protein complex (CORUM) enrichment was performed on proteins from neo N termini unique to patient or host. Complexes and pathways enriched in neo N termini with increased abundance in patients, were linked to protein pre-processing and maturation, as well as antigen processing and presentation (Fig. 5F). The sequence context for these neo N termini showed P1, P1’ and P2’ as the main specificity conveying positions (Fig. 5G) and multiple proteases, including matrix metalloproteinases, caspases, catalases, and granzymes have previously been shown to generate the observed termini (Fig. 5H). 12 neo N termini were downregulated in patients and showed a marked difference in their sequence composition from upregulated termini (Fig. 5I). Their corresponding proteins were enriched in the Histone H3.3 complex (CORUM, Benjamini Hochberg FDR = 0.032, enrichment factor = 101.62) and no matching protease cut-site had been reported in TopFIND.

In our smaller cohort of two T-ALL patients (blast count > 80%) and corresponding PDXs, 194 neo N termini were dysregulated (189 upregulated, and 9 downregulated) between patients and PDXs (Figs. 6A and 6B). Enrichment analysis showed similar processes and complexes associated with protein processing and maturation (Fig. 6C), comparable sequence motifs and a similar protease map (Figs. 6D-F), as was detected for B-ALL patients.

Proteases can cleave other proteases, and protease inhibitors, thereby indirectly regulating the cleavage of substrates of other proteases *in vivo*^37^. To explore protease interactions, we used PathFINDer^36^ to associate protein-level protease changes in leukemic models to direct or indirect cleavage resulting in observed N termini. Cathepsin S (CTSS) abundance was markedly lower or absent in PDX when compared to paired patient samples. In addition to the direct CTSS substrates, RPL7 and RAN, PathFINDer analysis linked neo N termini altered between T-ALL patients and PDX to potential downstream effects of CTSS activity (Fig. 6G). The first nine amino acids of Cystatin B (CSTB) are required for effective inhibition of Cathepsin B (CTSB) and Cathepsin L (CTSL) ^38^. Cleavage of Cystatin B by elevated Cathepsin S at position 9 could lead to increased activity of CTSB and CTSBL, and subsequent increase in neo termini of their substrates. Known products of CTSB and CTSL processing were present in patients and absent or reduced in xenografts, suggesting an activity-linked cleavage pattern for Cathepsin S in T-ALL patients. Proteins directly or indirectly linked to Cathepsin S *in vivo* proteolytic activity had functional roles in apoptotic signaling (GO:0072594 - ENO1, YWHAZ), RNA metabolism (R-HSA-8953854 - RAN, RPL7, RPL4, RPL13, YWHAZ) and protein folding (PPIA). A similar protease network was associated with neo N termini alterations in B-ALL (Supplementary Fig. 9D).

Although the vast majority of proteolytic processing events did not significantly differ between patient and PDX, a subset of the N terminome was significantly altered in PDX. The associated processes and protease-substrate relationships indicated differences in proliferation and apoptosis, further supporting the molecular differences identified at the protein level. The phosphorylation sites covered in this study did not significantly change with the transition from the patient to the mouse xenograft.

### Relapse specific changes in patients are retained in PDX

Our proteomic profiles could distinguish pediatric B-ALL and T-ALL subtypes at the protein (Fig. 1C), phosphoprotein, and N termini level (Fig. 2). We investigated if specific changes between diagnostic and relapse disease timepoints (Fig. 1A) were retained in xenograft leukemia (unpaired Student’s t-test FDR < 0.05). The profile of 108 proteins with significant abundance changes between B-ALL diagnostic and relapse patients was highly correlated between patient and matched xenograft (Fig. 7A). This included known proteins previously associated with leukemogenesis, including CD44 and STAT1 (Fig. 7B). Abundance differences at the neo N termini level were also largely comparable in patients and paired PDXs for 56 N termini that distinguished diagnosis and relapse conditions (Fig. 7C). N termini from leukemia-associated proteins such as PDK3 and LAT2 (Fig 7D) were elevated in relapse cases. Relapse associated changes in phosphosite levels were recapitulated in xenografts but showed lower conservation between matched pairs (Fig. 7E). Overall this shows that proteome and PTM profiles that distinguish diagnostic from relapsed childhood ALL are predominantly modeled in xenograft leukemias.

### PDXs recapitulate gene expression changes in patients

Some genetic changes in childhood B- and T-ALL leukemia have been linked to patient phenotype^1,3,7,39^. To determine how gene alterations in pediatric ALL are presented at the proteome level in patients and in patient-derived xenografts, we examined the profile of 202 pediatric cancer-related genes in the Oncomine Childhood Cancer Research Assay (OCCRA) targeted next-generation sequencing panel^32^. The recently reported OCCRA gene panel includes single nucleotide variation combinations, copy number variation and gene fusion combinations, that together proved more sensitive for the detection of childhood cancers as against adult-based panels. A substantial number of proteins from the OCCRA set of 201 genes were quantified in our sample cohort: 109 based on total protein abundance; 47 phosphoproteins, and N Termini matching to 34 proteins (Fig. 8A). Spearmans correlation scores, showed a strong similarity between total protein levels in patients and corresponding PDX (Fig. 8B). This was also the case for patients with more than one expanded xenograft mice (Supplementary Fig. 9E). Similarity scores were however lower for T-ALL patients with reduced bone marrow contribution and their corresponding xenografts (T-01-R1 and T-01-R1-PDXa/b, T-03-Dx and T-03-Dx-PDX respectively) (Fig. 8B). On average, the absolute Log2 protein levels in patients compared to PDX averaged below 1 in all pediatric ALL subtypes, indicating that the abundance profile of most OCCRA proteins were conserved in PDXs (Fig. 8C). Proteins KMT2A, KMT2D and GNA11 were the most variable proteins. Well-known pediatric cancer-relevant proteins are largely unchanged in primary and xenograft leukemia models, except for few proteins, in particular lysine methyltransferases 2A and 2D.

## Discussion

Patient-derived xenografts provide a vital means to simulate human diseases, understand disease biology, and most importantly, to develop therapeutic approaches. Comparative studies on patient and xenografted leukemia have established that genetic and epigenetic abnormalities associated with pediatric B- and T-cell leukemias are recapitulated in PDX models^7–11^. Proteins, in principle, drive and maintain cellular homeostasis in physiological and pathological conditions. Their functional relevance sets them apart as valuable indicators of disease phenotypes, and as therapeutic targets. It is therefore important, and of due time, to determine how accurately protein functional abnormalities in pediatric leukemia patients are reflected in PDX models. Such knowledge is of particular importance to determine if PDX models can be used to study response to treatments targeting signaling pathways in preclinical studies and if a particular model is suitable to study select aspects of cancer biology. Our results show that the protein landscape (proteome) of pediatric B- and T-cell leukemias is mostly retained in PDX mice with the exception of immune response and cell cycle pathways. The phospho- and proteolytic PTM landscapes of patient leukemia cells are largely retained in a xenograft environment but show a lower overall correlation than the more stable proteome.

While our ALL clinical and PDX cohort is insufficient for robust discovery of candidate disease markers, it offers for the first time, a global molecular view of disease-driven changes in the proteome of pediatric ALL and evaluates how this is recapitulated in the commonly used pediatric ALL xenograft NSG mouse model. Increased blast cell population in the bone marrow of patients with B- and T-cell leukemias results in a protein composition that is distinct from the joint proteome of bone marrow mononuclear cells isolated from non-leukemic patients. The homogeneity of the analyzed mononuclear cell population is a major contributing factor in this as evident from the near normal (non-leukemic) proteome profile found in leukemic patients with low leukemic bone marrow involvement of 2 and 10% respectively. Patient leukemic cells from diagnosis and relapse conditions stably retained their protein-level characteristics after xenotransplantation in immunodeficient mice. This was also evident in biological replicates where blast cells from the same patient were transplanted into two different mice. In addition to this, the PDX model capably propagated leukemias from patients with minimal bone marrow contribution, and more importantly retained a leukemia protein landscape similar to other patients and PDXs. Post-translational protein modification by phosphorylation and proteolysis are amongst multiple modification events that expand the functional roles of a protein. Proteolytic processing assessed by N terminome profiling, and phosphorylation profiles of patients are clearly distinct from non-leukemic bone marrow mononuclear cells and show clear overall correlation with PDXs. Patient specific differences in post-translational modifications are not always recapitulated in the matched xenograft. As this study was limited to one specimen per patient it was not possible to determine how robust these differences reflect the individual patient’s disease and if the limited correlation of matched patient / PDX pairs entails robust differences in treatment relevant pathways. Establishing a cohort with longitudinal sampling is particularly challenging as repeat biopsy collection from young patients is ethically problematic. Thus, we could not conclusively determine if xenografts are adequate to model patient specific differences mediated by proteolysis or phosphorylation. This would warrant targeted perturbation with compounds of interest as well as comprehensive longitudinal sample profiling. The study does however provide strong support for xenografts as accurate model of pediatric acute leukemias. The data confirms that disease-related, steady-state phosphorylation profiles are maintained in PDX and patient host.

Pediatric leukemias have diverse genomic aberrations ^1,3,39^, known to be replicated in xenografts^7,40^. With proteomics, we confirmed that structural defects and mutations in patients are faithfully recapitulated in their xenografts. An overview of the 219 protein abundance features associated to 883 protein-coding genes on chromosome X indicated that patients with trisomy X had an abnormal protein profile, as did their paired PDXs. Additionally, proteomics and targeted next-generation sequencing on 6 paired samples confirmed our findings. Specifically, protein absence and depletion accompanied copy number losses in *CDKN2A* and *RB1* respectively. Of note, the protein profile of 84% of pediatric cancer-related genes (Fig 8C) validated from extensive genomic and transcriptomic characterizations ^3,7,32,39,41^ was essentially preserved in xenograft cells after expansion in mice.

Other novel findings in this study exposed deficient or missing components in the PDX model that may compromise its capacity to replicate specific processes and pathways in patients. Genomic and epigenomic characterization of T-cell leukemia PDX models did show reduced immune/defense responses and limited stimulation of cytokine response in xenografts compared to their paired primary leukemias ^11^. This was attributed to increased chromatin condensation of gene regulatory elements involved in immune function and regulation of cytokine elements. Our study showed that such compromised immune and defense responses are also evident at the proteome level of T-ALL and similarly found in B-ALL. In B-ALL, this was restricted to the loss of vital complement proteins, and cell-surface glycoproteins. While T-ALLs particularly lacked major proteins involved in the activation of the T-cell receptor and in MHC Class II -mediated antigen presentation. Our data profiled the cellular levels of 180 proteins linked to cytokine-mediated signalling pathways, cytokine production, and cytokine metabolic processes. The majority of these proteins were stable between PDX and patients. However, five proteins were markedly deficient or missing in both B- and T-cell xenograft leukemias (Supplementary Fig. 10A). Proteins S100A8, S100A9, S100A12 are important for cytokine production, while the secreted glycoprotein, CHI3L is a Th2 promoting cytokine. Our data also identified increased abundance of proteins linked to cell cycle and mitosis in PDXs over patients suggesting increased proliferation in xenografted leukemia cells. This is in line with genome-wide DNA analysis on paired T-ALL patients and PDXs which suggested that increased proliferation contributes to the selective advantage of engrafted cells^7^. Increase in cell cycle and proliferation markers may also be explained by a lack of spatial constraints for xenografted cells localized to the spleen relative to patient cells restricted by the bone marrow confinement. The presence of a greater fraction of proliferating blasts in PDX spleen than in patient bone marrow is supported by the observation that xenografted cells continue to expand indefinitely in the murine spleen.

This study provides initial insights into the fate of ALL-related post-translational modifications in xenografted blasts. T-ALL and B-ALL xenografts retain a ‘leukemic’ N terminome, similar to patients with high leukemic burden, and distinguishable from non-leukemic patients and patients with lower leukemic bone marrow burden. Neo N termini patterns from limited proteolysis act as fingerprints to identify active proteases and their corresponding cleavage sites in different cellular conditions and locations^24^. Indeed, these patterns revealed non-normal enriched proteolytic activities in pediatric ALL cell lines, patients and PDXs. They further suggested that blast cells loose CASP1 proteolytic activities after transplantation in NSG mice, while gaining additional processing functions for ELNE (Supplementary Fig. 10B). Constrained CASP1 activities in PDXs supports the reduced presence of cytokines in blast cells xenografted in immunocompromised NSG mice. The unique cleavage pattern of MCM5 in patients and corresponding PDXs further substantiates the existence of proteolytic fingerprints that are indistinguishable at the protein level (Supplementary Fig. 10C). MCM5 mature protein N term was consistently detected in PDXs, while the neo N termini products from MCM5 cleavage by HTRA2/GZMM proteases were principally absent in PDXs. Unlike MCM5 N termini, protein amount in patients and matched PDXs was mostly stable.

The absence of functional immune response processes in xenografted NSG mice is most striking at the proteome level, and points to the absence of the immune component in NSG mice, and the inability of related murine proteins to substitute their human counterparts^42^. The results reported in this study further indicate absence as well as limited abundance of specific immune-regulatory proteins in the immunocompromised murine environment seem to dampen the stimulation of immune pathways in xenografts. Furthermore, some protein complexes and interactions are possibly disrupted in engrafted leukemia. Our data show perturbed abundance of interacting complement factors and human leukocyte antigen MHC class II cell surface receptors in transplanted models. Likewise, multi-complex protein subunits with major roles in RNA processing, ribosome biogenesis, and protein maturation are altered following xenografting. These discrepancies in proteome representation should be considered when PDX models are utilized in preclinical investigation.

In conclusion, our study shows for the first time that pediatric leukemia cells principally maintain their protein abundance and modification pattern when xenografted into immunocompromised NSG mice. Total protein and protein modification landscapes appear differently affected by the host, leukemia type and patient of origin. Phosphorylation and proteolysis are largely correlated between patient and xenograft but less robustly recapitulate patient and xenograft pairing. These differences underscore the need to characterize not only protein abundance but also key post-translational modifications and we here provide a proof of concept, that the required combined proteome and PTM analyses can be performed on clinical specimen consisting of as little as 600,000 cells.

For the most part the proteome and PTM landscapes persist following transplantation of human B-ALL and T-ALL blasts into NSG mouse. PDX models remain the most fitting proxy for investigation of subtype and patient-specific disease biology, and evaluation of new therapeutic approaches.

## Materials and Methods

### Experimental Samples

#### Cell lines

B-ALL cell lines 380 (ACC 39), 697 (ACC 42) and T ALL cell lines DND-41 (ACC 525), PEER (ACC 6) were purchased from DSMZ (Braunschweig, Germany). Cell lines were cultured in RPMI-1640 media supplemented with 10% heat-inactivated fetal bovine serum (FBS) and 2 mM L-Glutamine (Gibco, Grand Island, NY) and maintained at 37 °C in 5% CO_2_.

#### Patient bone marrow and peripheral blood samples

patient samples were collected with informed consent by Biobank staff during routine clinical care at BC Children’s Hospital (BCCH). Sample collection and experiments were performed as approved by the University of British Columbia Children & Women’s Research Ethics, and conformed with standards defined in the WMA Department of Helsinki and the Department of Health and Human Services Belmont Report. Mononuclear cells containing leukemic blasts were isolated by Ficoll-Paque PLUS density centrifugation, viably frozen and preserved. Aliquots of patient samples, and patient clinical information were de-identified prior to release for this study.

As part of routine clinical care, patient samples were immunophenotyped by trained staff at the clinical hematopathology laboratory using established ALL subtype-specific 10-colour flow cytometry panels according to clinical standard operating procedures. Patient bone marrow morphology was routinely assessed by hematopathologists and clinical cytogeneticists performed cytogenetic studies on leukemia samples.

Clinical cohort comprised of 8 patients with B-cell ALL (4=diagnosis, 4=relapse), 3 patients with T-cell ALL (2=diagnosis, 1=relapse), one patient with T-cell lymphoblastic lymphoma (T-LBL) and 2 patients with no detected leukemic blasts (normal or non-leukemia) (Table 1). Patient age ranged from 3.9 to 17.7 months. The mean age of patients was evenly distributed between gender (females = 10.3, males = 10.7 months). And the extent of leukemic blast infiltration to the bone marrow of patients was between 2 and 97%. Patient N-01 had GATA2 mutation, and patient N-02 had Thrombocytopenia. Mutation of the gene encoding the hematopoietic transcription factor *GATA*2 predisposes children to diverse hematologic malignancies^43^. Individuals with this mutation have a broad spectrum of clinical manifestations ranging from haematologically normal to mononuclear cytopenia, and immune dysfunction^43,44^. Carriers are clinically monitored for hematologic disorders or other life-threatening complications. Our inclusion of this patient in our study cohort is justified for the following reasons. First, history of patient follow-up (until 73 months after initial biopsy) showed no symptoms for any hematologic malignancies. No predisposing cytogenetic abnormalities, including monosomy 7 were detected in this individual. Also, despite germ line GATA2 mutation, the patient till date lacks symptoms of primary myelodysplastic syndrome (MDS). Patient N02 also remained asymptomatic as at last clinical observation (61 months post diagnosis).

#### Patient-derived xenografts

Primary ALL cells engraftment in mice was performed in accordance with an Institutional Animal Care and Use Committee-approved protocol (A15-0187). Briefly, viable mononuclear cells (MNCs) in sterile PBS were injected into the tail vein of 6- to 10-week old NOD.Cg-Prkdc^scid^/IL2rg^tm1Wjl^/SzJ (NSG) mice initially purchased from The Jackson Laboratory and bred and maintained in-house under pathogen-free conditions. Mice were euthanized at onset of overt leukemia which was determined by flowcytometry. Mononuclear cells isolated from harvested bone marrow and spleen using standard procedure were viably frozen and stored.

### Targeted Next Generation Sequencing

Amplicon-based sequencing and variant determination was performed as described elsewhere^32^. Briefly, DNA and RNA extraction were performed using the RecoverAll (Thermo Fisher Scientific) and Allprep (Qiagen) workflow package respectively. Library preparation and targeted sequencing on an Ion Chef and Ion Torrent S5 platforms (Thermo Fisher Scientific) followed manufacturers’ protocols. The Oncomine Childhood Cancer Research Assay (OCCRA) comprises 2,031 unique DNA-based amplicons to detect SNVs, and CNVs, and 1701 RNA-based amplicons to detect unique fusion or structural variants. The average read depth for the OCCRA panel was 8×10^5^ – 1×10^6^.

SNVs, including those in pediatric cancer driver genes, non-driver genes, variants of undetermined significance and benign/likely benign variants, were retrieved with Ion Reporter software (version 5.2). Copy number measurements were retrieved with Ion Reporter software (version 5.2) for genes with >5 probes, including those that were validated for copy number gains.

### Preparation of samples for mass spectrometry-based analysis

Suspension cells cultured in T75 flasks were passaged up to five times. Biological triplicates were cultured separately. Cells harvested at greater than 90% confluency were washed twice with 1% PBS (Gibco, Grand Island, NY) and pelleted by centrifugation.

Mononuclear cells isolated from patient bone marrow and blood, and from mice spleen, were quickly thawed, washed twice with 1X PBS, and harvested by gentle centrifugation. Cell viability performed on a 10 μl aliquot was above 70%.

Except otherwise stated, reagents were purchased from Sigma Aldrich (St. Louis, Missouri, United States). Cells were lysed in buffer containing 1% SDS (Fisher BioReagents, Pittsburgh, Pennsylvania, United States), 1X Pierce protease inhibitor (Thermo Fisher Scientific, Waltham, Massachusetts, United States) in 50 mM HEPES (pH 8.5), followed by 5 min incubation at 95 °C and 5 min at 0 ^o^C, on ice. The supernatant containing proteins was collected after centrifugation and incubated in benzonase (EMD Millipore/Novagen, Country) at 37 °C for 30 min to shear chromatin. Protein amount was determined by Bicinchoninic acid (BCA) assay.

#### Proteome and phosphoproteome

For each cell line, patient and PDX sample, 60 μg protein was reduced with 10 mM Tris-(2-carboxyethyl)-phosphine (TCEP) (37 °C, 1 hr), followed by alkylation with 40 mM Chloroacetamide (CAA) (45 min in the dark) and quenched in 40 mM TCEP for 10 mins at room temperature. Reduced lysate was further processed with single-pot solid-phase-enhanced (SP3) bead technique^45^ using hydrophilic and hydrophobic Sera-Mag Speed Beads (GE Life Sciences, Issaquah, Washington, United States). Proteins were bound to paramagnetic beads with 100% ethanol (80% v/v), incubated for 18 min at room temperature, and washed twice with 90% ethanol using magnetic isolation. Beads were then resuspended in 50 μl 200 mM HEPES, pH 8.0, and incubated with sequencing-grade trypsin (Promega Madison, Wisconsin, United States) at 1:20 protein ratio for sixteen hours at 37 °C, and afterwards acidified to pH 3-4 with formic acid. Peptide digests were washed on a Nest Group Inc. C18 spin columns with 2% acetonitrile in 0.1% formic acid, eluted with 80% acetonitrile in 0.1% FA and dried in a Speed Vac.

Dried samples were resuspended in 0.1% TFA in 80% acetonitrile and phosphorylated peptides were purified by immobilized metal affinity chromatography (IMAC) using Fe-NTA MagBeads (Cube Biotech, Monheim, Germany). Sample solution was added to beads washed with 0.1% TFA in 80% acetonitrile, and incubated in a thermomixer at room temperature for 30 min. Beads were magnetically isolated to recover the non-phosphorylated peptide fraction in the supernatant. Following two wash steps in sample resuspension buffer, phosphorylated peptides were eluted from Fe-NTA beads using 1% ammonia, and immediately acidified to pH 3 with formic acid.

Peptide solutions were cleaned with The Nest Group Inc. C18 spin columns by washing with 0.1% TFA, and eluted from the column with 0.1% FA in 50% acetonitrile. After drying in a Speed Vac, samples were resolubilized in 0.1% FA for mass spectrometry analysis.

#### N-termini peptide enrichment using HUNTER technique

For each sample, 30 μg protein was reduced with 10 mM dithiothreitol (DTT) (37 °C, 1 hr), followed by alkylation with 50 mM chloroacetamide (CAA) (45 min in the dark) and quenched in 50 mM DTT for 10 mins at room temperature. Reduced lysate were further processed with paramagnetic single-pot solid-phase-enhanced (SP3) beads^45^ as described earlier, resuspended in 30 μl 200 mM HEPES, pH 8.0, and incubated with sequencing-grade trypsin (Promega Madison, Wisconsin, United States) at 1:30 protein ratio for sixteen hours at 37 °C. Peptide labeling and enrichment was performed as reported elsewhere^25^. Briefly, proteins were dimethylated by incubating lysates in 30 mM formadehyde and 15 mM sodium cyanoborohydride for 1 hr at 37 °C, pH 6-7, after which same amounts of formaldehyde and sodium cyanoborohydride was added and incubation conditions were maintained for another 1 hr. The reaction was quenched by adding 600 mM of Tris (pH 6.8) and incubating for 3 hr at 37 °C. After a second round of bead binding in same volume of SP3 beads (10 μl) and 100% ethanol (80% v/v), beads were washed three times with 90% acetonitrile and briefly air-dryed. Beads were then resuspended in 30 μl 200 mM HEPES, pH 8.0, and incubated with sequencing-grade trypsin (Promega Madison, Wisconsin, United States) at 1:30 protein ratio for sixteen hours at 37 °C. Negative selection of N-termini peptide digests was done using undecanal tagging. 100% ethanol was added to a final concentration of 40% v/v, followed by undecanal (20 μg undecanal per 1 μg peptide), and 30 mM sodium cyanoborohydride, pH 7-8. After 1 hr incubation at 37 °C, peptides were acidified to pH 3 with 0.5% TFA in 40% ethanol. To get rid of undecanal, samples were cleaned with Nest Group C18 midi spin columns: cleaned with methanol and conditioned with 0.1% TFA in 40% ethanol. Sample flow-through was collected and dried in a Speed Vac. Dried samples were resuspended in 0.1% TFA in water for a second C18 clean up, washed with 0.1% TFA in 2% acetonitrile, and eluted in 0.1% FA in 40% ethanol. After drying in a Speed Vac, samples were resolubilized in 8 μl of 0.1% FA in HPLC water, prior to mass spectrometry analysis.

### High pH Reversed Phase Fractionation

For non-phosphorylated peptides, 1 μg from each sample was combined and fractionated. A pool of phospho-enriched peptides was generated by combining 1 ul of phosphopeptides from each sample. HUNTER-enriched N term peptides from each sample was combined (1 ul per sample) and fractionated. Fractionation was performed on a Kinetic EVO C18 column (2.1 mm x 150 mm, 1.7 µm core shell, 100Å pore size, Phenomenex) connected to an Agilent 1100 HPLC system equipped with a diode array detector (254, 260, and 280 nm). A flow rate of 0.2 ml per minute was maintained on a 60 min gradient using mobile phase A (10mM ammonium bicarbonate, pH 8, Fisher Scientific, cat. no. BP2413-500). Elution was with mobile phase B (acetonitrile, Sigma-Aldrich, cat. no. 34998-4L) from 3% to 35%. Peptide fractions were collected each minute across the elution window. A total of 48 fractions were combined to a final set of 12 (e.g fraction 1 + 13 + 25 + 37 as final fraction 1), and dried in a SpeedVac centrifuge. Peptides were resuspended in 0.1% FA in water (SC235291, Thermo Scientific) prior to mass spectrometry analysis.

Preceding mass spectrometry analysis, lyophilized peptides samples were resolubilized in 0.1% formic acid, spiked with index Retention Time peptides (iRT, Biognosys, Switzerland).

### Mass Spectrometry (LC-MSMS) Analysis of Samples

#### Data Dependent Acquisition for Spectral Library Generation

Peptide solutions were analyzed using a Thermo Scientific Easy-spray PepMap™RSLC C18 column (75 μm x 50 cm, 2 μm, 100Å; ES803), maintained at 50 °C on an Easy-nLC 1200 connected to a Q Exactive HF mass spectrometer (Thermo Scientific). Peptides were separated over a 2 h gradient consisting of Buffer A (0.1% FA in 2% acetonitrile) and 3 to 32% Buffer B (0.1% FA in 95% acetonitrile) at 250 nL/min. MS acquisition was performed with full scan settings between 350 and 1600 m/z, resolution of 120000, AGC target of 1 e6, and Maximum IT of 246 ms. Stepped collision energy (NCE) was 28. MS2 scan settings were as follows: isolation window of 1.4 m/z, AGC target of 2 e5, maximum IT of 118 ms, at resolution of 60,000 and dynamic exclusion of 15.0 s.

#### Data Independent Acquisition

One micro gram per sample of resolubilized peptides were analyzed using a self-packed analytical PicoFrit column, 75 μm x 35 cm length (New Objective, Woburn, Massachusetts, US), packed with ReproSil-Pur 120A C18-AQ 3 μm (Dr. Maisch GmbH, Ammerbuch, Germany) at 50 °C Germany. Peptide separation was performed on an EASY-nLC 1200 connected to a Q Exactive HF mass spectrometer (Thermo Scientific) using a 1 hr gradient consisting of Buffer A (0.1% FA in 2% acetonitrile) and 1 to 34% Buffer B (0.1% FA in 95% acetonitrile) at 300 nL/min.

Approximately 1 μg of whole protein per sample was injected, and the DIA method (adapted from Bruderer et al.,^46^), consisted of a MS1 scan from 300 to 1650 *m/z* (AGC target of 3e6 or 60 ms injection time), and resolution of 120,000. DIA segment spectra was acquired with a twenty-four-variable window format, (AGC target 3e6, resolution 30,000, auto for injection time), and stepped collision energy of 10% at 25%. Samples were injected in duplicates, except for phosphopeptide samples which had single injections. The combined sample pool was injected for DIA analysis in between the sample acquisition to monitor instrument performance. For N-Terminome analysis, a 10 variable window format was used with resolution of 60,000, AGC target of 3e6, auto for intejction time, and collision energy of 28.

At each stage of sample preparation and MS acquisition, - whole protein, phosphoprotein and N-termini, samples were randomized and blocked accordingly.

#### DIA Data Analysis

DIA data was analyzed with Spectronaut Pulsar X (version 12.0.20491.3.15243, Jocelyn from Biognosys, Schlieren, Switzerland)^47^. First, a spectral library was generated for each experiment – whole proteome, phosphoproteome, N-Terminome, in Spectronaut Pulsar by searching DIA sample raw files together with the 12 DDA files from high-pH fractionation. The default Spectronaut Pulsar search settings were applied and included: digest type as Specific for Trypsin/P enzyme cleavage with two missed cleavages, carbamidomethyl (C) as fixed modification, Acetyl (Protein N-term) and Oxidation (M) as variable modifications. Same settings were maintained for phosphopeptide analysis, with Phosphho (STY) as additional variable modification. N-Termini settings were as follows: Semi-specific (free N-terminus) digest by Arg-C, carbamidomethyl (C) and DimethylLys0 as fixed modifications, while variable modifications consisted of Acetyl (N-term), DimethylNter0, Gln->pyro-Glu, Glu->pyro-Glu, and oxidation (M). Default library generation parameters were used at a minimum Rsquare of 0.8 for the iRT peptides. The ideal mass tolerance for MS1 and MS2 searches were determined based on extensive mass calibration (Dynamic setting) and a correction factor of 1 was applied at both levels. A false discovery rate threshold of 0.01 was maintained for peptide spectra matching, peptide and protein identification. Spectra was matched against the Uniprot-Swissprot human protein fasta database (30-05-2019) with 20,874 entries. Fragment ions with amino acid length greater than 3, and *m/z* range of 300 to 1800 were included in the spectral library. The best three to six fragments were selected per peptide. The resulting spectral library contained fragment annotation and normalized retention times for each experiment, as follows (1) Whole proteome: 122,510 precursors matched to 94,103 peptides and 7,969 protein groups. (2) Phosphoproteome: 9,550 precursors matched to 8,199 modified peptides and 1,836 protein groups. (3) N Terminome: 6,910 precursors matched to 5,282 modified peptides and 2,080 protein groups.

Each spectral library was used for targeted analysis of DIA data using the default Spectronaut settings^46,47^. In brief, MS1 and MS2 tolerance strategy were ‘Dynamic’ with a correction factor of 1. Similar setting was maintained for the retention time window for the extracted ion chromatogram. For calibration of MS runs precision iRT was activated, with local (non-linear) regression. Feature identification was based on the ‘mutated’ decoy method, with ‘dynamic’ strategy and library size fraction of 0.1. Minor (peptide) grouping for quantification was by ‘stripped sequence’ for whole protome, and by ‘modified sequence’ for both phosphoproteome and N-Terminome analysis. Precursor and protein false discovery rates were 1% respectively.

### Statistical analysis and interpretation of MS data

Spectronaut default output files from whole proteome (Supplementary Dataset 1 and 2) and N-Terminome (Supplementary Dataset 3 and 4) experiments were exported for data analysis and interpretation. Each experiment data was filtered to remove entries with less than 8 values, and retain samples with duplicate injections that passed quality control assessments. N-Terminome data was further processed to retain only Dimethyl (N-Term) and Acetyl (N-Term) modifications. Median centering normalization was performed on the filtered data (whole proteome = 40 of 45 samples, N-Terminome = 42 of 43 samples).

Precursor information from DIA analysis on phophopeptides were exported from Spectronaut and processed with the Peptide Collapse PlugIn (version v1.4.1) ^48^ in Perseus (version 1.6.2.2)^49^ using the recently reported peptide-centric, PTM-specific workflow for high-confidence site localization and quantification of phosphorylation sites^48^. The following settings were used: collapse level = Target PTM site-level, probability column type = EG.PTMLocalizationProbabilities (SN), localization cut-off = 0.75, Target PTM = [Phospho (STY)], Variable PTMs = [Acetyl (Protein N-term)] and [Oxidation (M)], aggregation type = linear modeling based. The resulting quantified data (Supplementary Dataset 5 and 6) were used for further analysis

Quantified, filtered datasets were normalized (verified by frequency distribution plots). Unsupervised t-distributed stochastic neighbor embedding (t-SNE) was done on Log_2_ transformed protein intensities with the following parameters: n_components=2, perplexity=10, n_iter=10000, random_state=1). Missing values in data were imputed with values from lower tenth percentile of the whole data and down-shifted by a factor of 50, making sure the imputed values represent the lowest intensity of the data. Statistical analysis was also done with two-tailed Student’s t-test. And where applicable, data was adjusted for multiple comparisons.

Data processing, and figure plotting was done in R, python, Perseus, GraphPad Prism and BioVinci. Hierarchical clustering analysis was performed with ‘Euclidean’ distance and ‘Average’ linkage for row and column trees.

### Characterization of N-terminal peptides

Identified N-terminal peptides were processed using TopFINDer web program, which integrates available TAILS and COFRADIC N-terminome data and information from MEROPS database, to annotate known proteolytic processing events, and identify the characteristics of N-terminal peptides^34,36^. N-terminal peptides with no curated UniProt start site, no alternative start, no alternative translational start site, and P1 protease cut-site start at or after amino acid 5 were grouped as neo N termini. Protease network interactions were queried in PathFINDer^36,37^. Sequence pattern enquiry and graphical display of aligned consensus was performed using WebLogo (Version 3)^50,51^.

### Biological process and pathway enrichment

Protein pathway enrichment and protein network analysis was performed using the web-based portal for Metascape^30^. Representation and editing of network plots was done in Cytoscape (Version 3.7.1)^52^. Gene annotation and enrichment analysis on differentially regulated features was performed using the Fisher Exact Test option in Perseus.

## Acknowledgements

We thank patients and families, doctors and nurses, clinical and Biobank staff at the BC Children’s Hospital. This work was performed within the framework of the Better Response through AVatomics Evidence (BRAVE) initiative at BC Children’s Hospital directed by P.F.L., G.R.S., C.J.L. and C.M. and funded by the Michael Cuccione Foundation. This work was partially supported by grants from the BC Proteomics Network (to P.F.L), the Michael Cuccione Foundation and the BC Children’s Hospital Foundation (to P.F.L.). A.C.U., L.N. and E.K.E. were supported by fellowships from the Michael Cuccione Childhood Cancer Research Program or BC Children’s Hospital Research Institute. P.F.L. was supported by the Canada Research Chairs program and the Michael Smith Foundation for Health Research Scholar program.

## Author contributions

A.C.U., and P.F.L. designed the research; P.F.L., G.S.R., C.M. and C.J.L. initiated, designed and directed the BRAVE framework initiative; A.C.U. performed proteomic experiments with assistance from J.T., L.N., and S.S.W; G.S.R. and N.R. established and performed xenograft expansion; A.L. performed targeted next-generation sequencing; A.C.U., E.K.E., and T.S analyzed data; A.C.U. and P.F.L. wrote the manuscript with contribution from all authors.

