## Supplemental Figures and Tables for "PDX models reflect the proteome landscape of pediatric acute lymphoblastic leukemia but divert in select pathways"

Running title: Proteome stability in pediatric PDXs

### Supplementary Figures

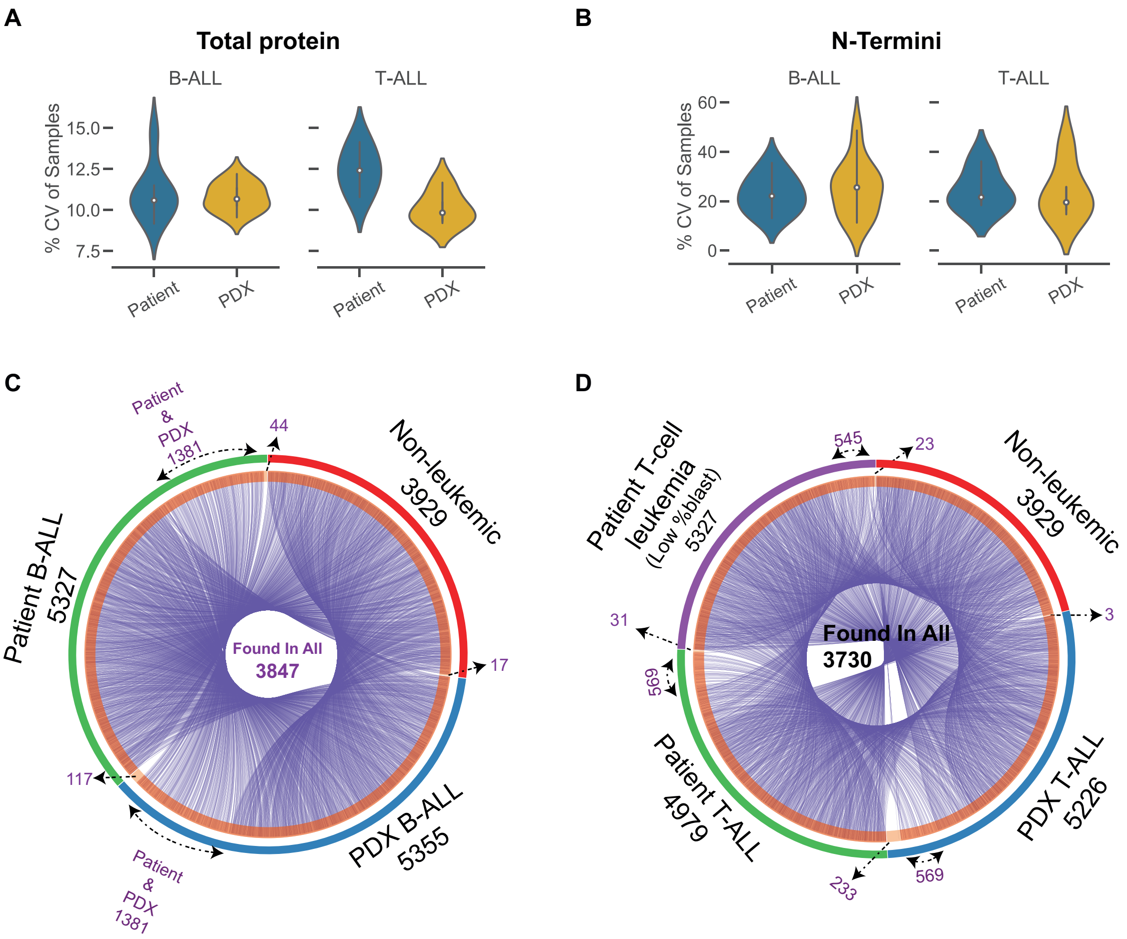

**Uzozie et al., Supplementary Figure 1**

**Supplementary Figure 1 - Experimental variation in proteomics data and protein-level overlap.**
A, B. The distribution of coefficient of variations (CVs) in patients and PDXs from B-ALL and T-ALL, determined for (A) total protein abundance (N = 5554) and (B) N termini abundance (N = 2832) calculated from replicate DIA sample measurements.
C, D. Commonality of protein features identified in ALL patients and PDX of (C) B-cell and (D) T-cell origin, and in non-leukemic samples.

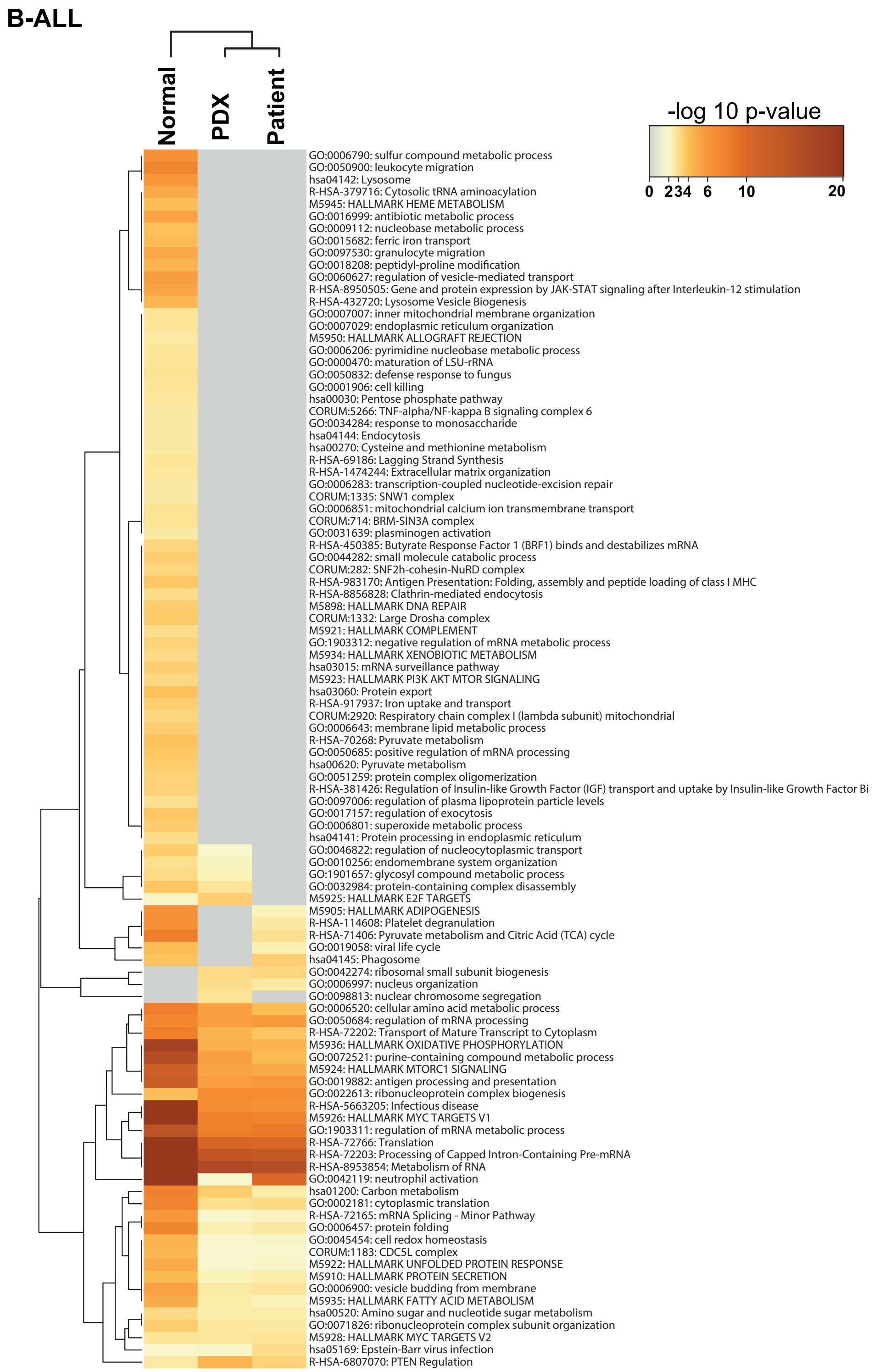

**Uzozie et al., Supplementary Figure 2**

**Supplementary Figure 2 - Biological pathways and processes enriched in non-leukemic, B-ALL patients and B-ALL xenografts.**
Heatmap of the Top 100 enriched terms in non-leukemic patients, B-ALL patients, and B-ALL xenografts. The colour scale of the heatmap is based on the p-value threshold. Enrichment analysis in Metascape is detailed in Methods.

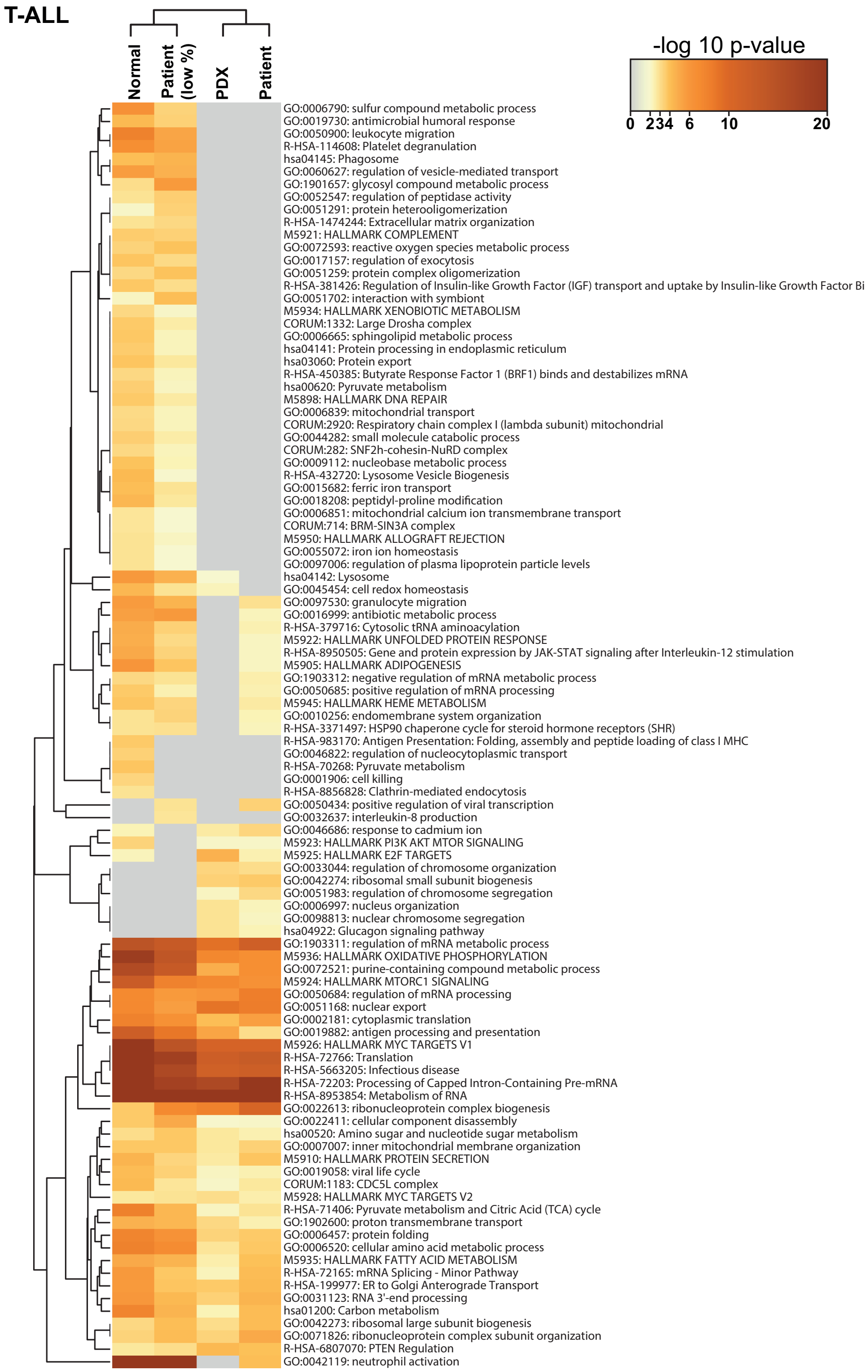

**Uzozie et al., Supplementary Figure 3**

**Supplementary Figure 3 - Biological pathways and processes enriched in non-leukemic, T-cell leukemia patients with low and high blast count, and T-cell leukemia PDXs.**
Heatmap of the Top 100 enriched terms in non-leukemic patients, T-cell leukemia patients with blast counts less than 10 percent, T-ALL patients with blast counts of 80% to 100%, and T-ALL xenografts. The colour scale of the heatmap is based on the p-value threshold. Enrichment analysis in Metascape is detailed in Methods.

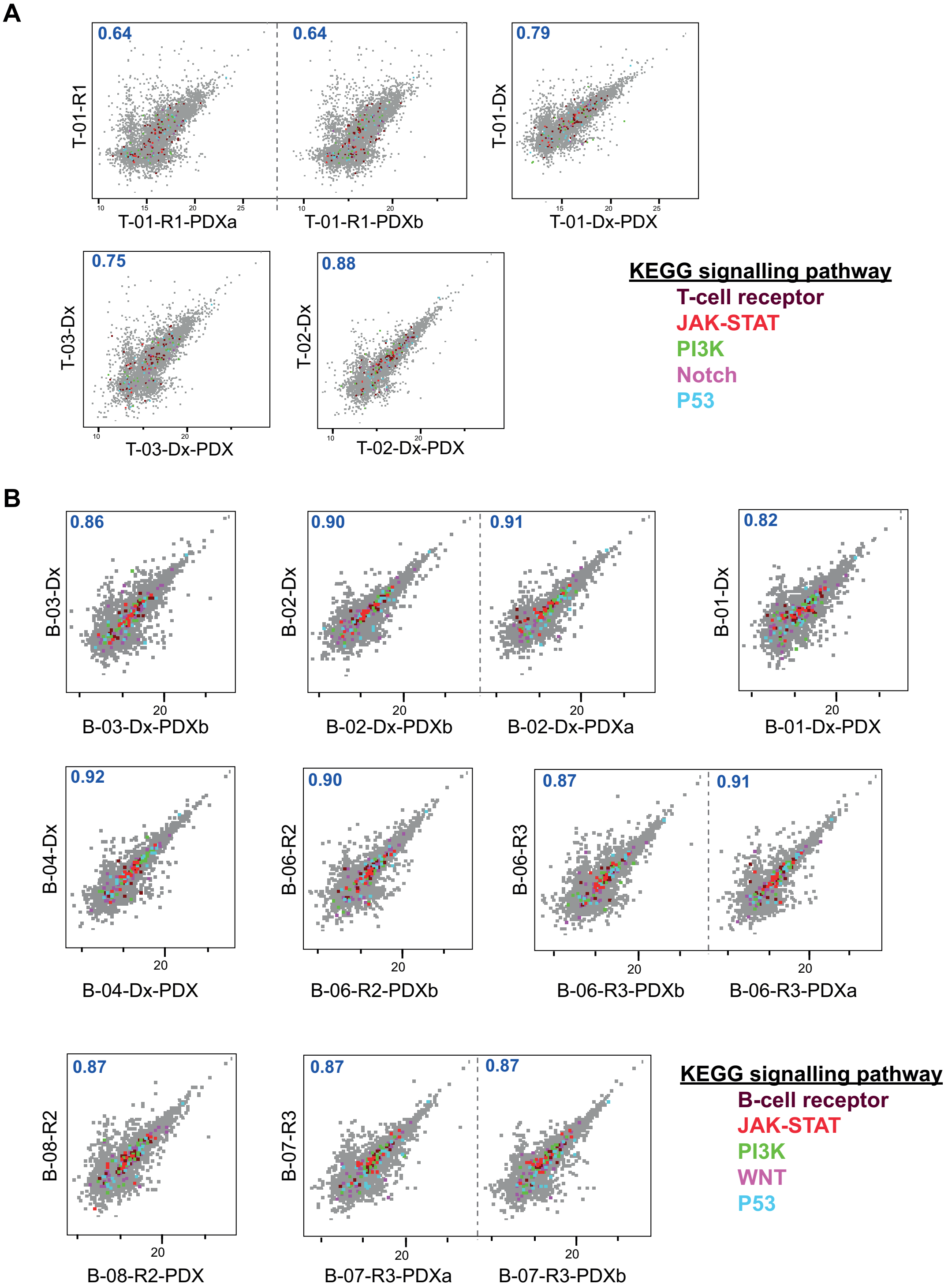

**Uzozie et al., Supplementary Figure 4**

**Supplementary Figure 4 - Protein-level correlation of paired patient and PDX samples.**
A. Multiple scatterplots on protein intensities (N = 5554) in matched primary and xenograft T-ALL samples.
B. Multiple scatterplots on protein intensities (N = 5554) in paired primary and xenograft B-ALL samples.
Spearman rank correlation score is highlighted in blue for each pair. Colour-coded points indicate proteins associated with KEGG signalling pathways affected by known gene mutations in ALL.

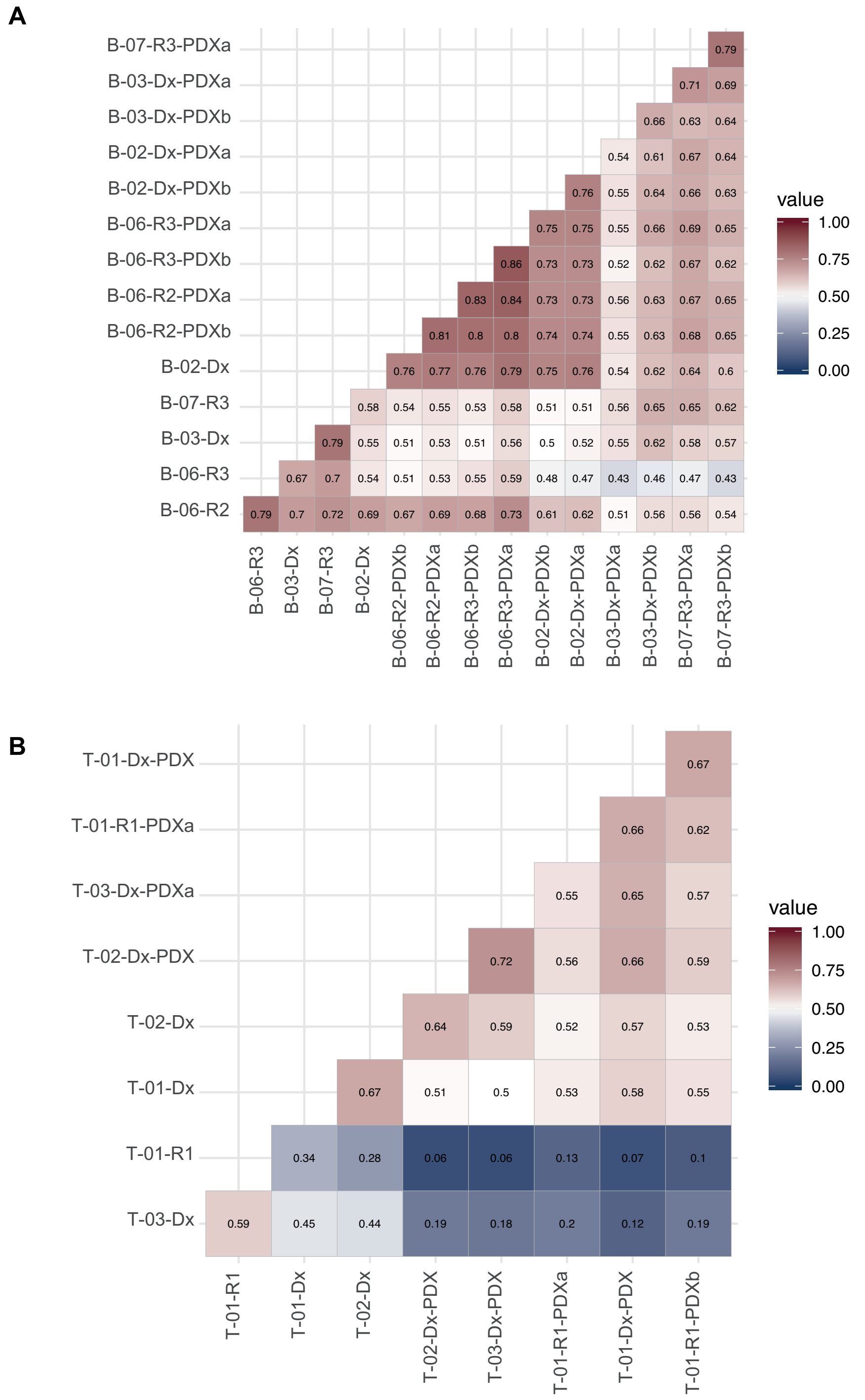

**Uzozie et al., Supplementary Figure 5**

**Supplementary Figure 5 - Correlation of N termini features in paired patient and PDXs.**
A, B. Correlation plots showing similarity in N termini abundance (N=2832) between (A) matched B-cell primary and xenograft leukemia and (B) paired T-cell primary and xenograft leukemia. Values inset are Spearman rank correlation score for each pair.

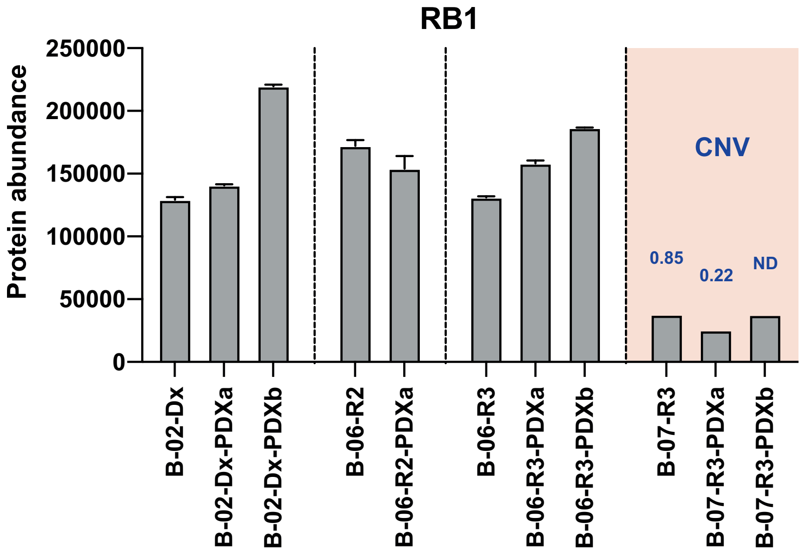

**Uzozie et al., Supplementary Figure 6**

**Supplementary Figure 6 - Consequence of *RB1* mutation on protein abundance.**
RB1 protein abundance in PDXs not measured by targeted sequencing (B-02-Dx-PDXa, B-06-R2-PDXa, B-06-R3-PDXa, B-07-R3-PDXb) are plotted together with samples analyzed by targeted sequencing. Similarity in RB1 level for both PDXs from patient B-07-R3 with *RB1* CNV of 0.0.85 is highlighted. ND means no sequencing done on sample. Bar plots show mean protein abundance from technical replicate DIA measurements and error bars represent the standard deviation from the mean.

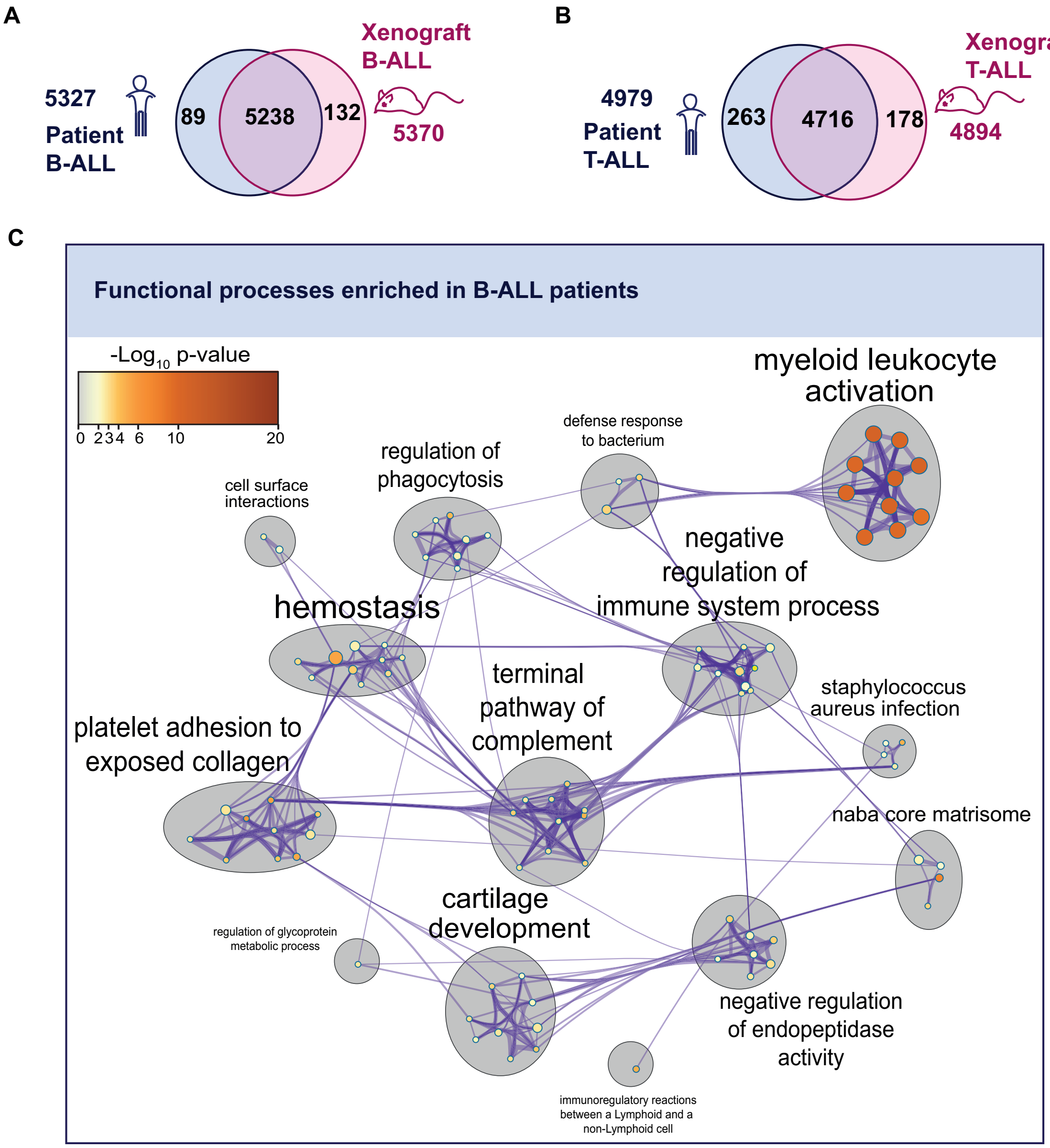

**Uzozie et al., Supplementary Figure 7**

**Supplementary Figure 7 - Biological processes that differ in human and PDX ALL.**
A. Overlap of proteins identified in patient and xenograft B-ALL.
B. Overlap of proteins quantified in patient and xenograft T-ALL, excluding identifications in samples from patients with minimal bone marrow blast involvement.
C. Network plot showing functional processes enriched in 89 proteins unique to patient B-ALL samples studied.
Data information: In (C), each circle represents a non-redundant cluster term. Circle size and colour are based on level of significance (-$Log_{10}$ p-value). Clusters comprise connected enriched terms, and the most representative category is highlighted (additional details in Methods).

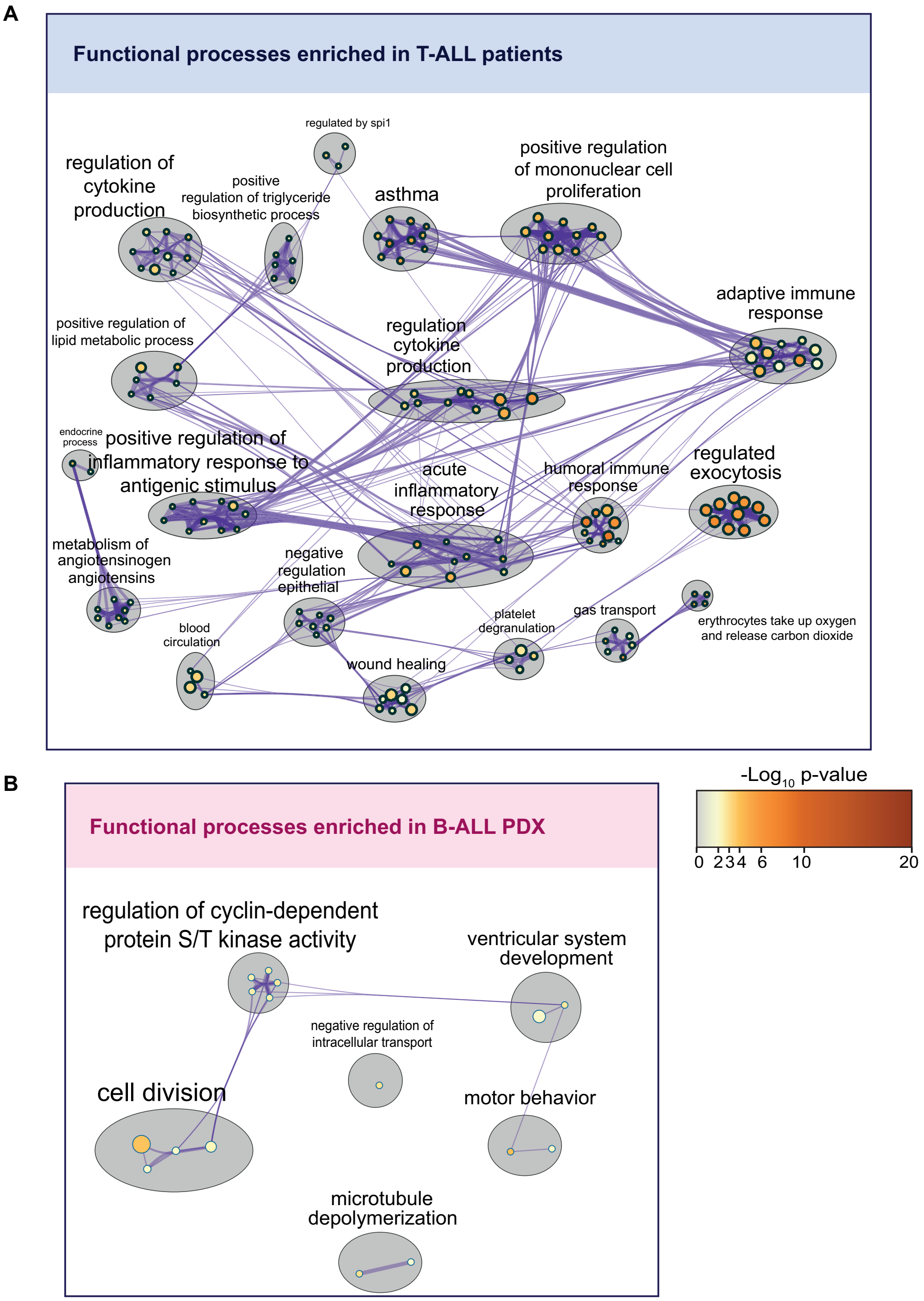

**Uzozie et al., Supplementary Figure 8**

**Supplementary Figure 8 - Biological processes that differ in human and PDX ALL.**
A. Network plot showing functional processes enriched in 263 proteins unique to patient T-ALL samples studied.
B. Network plot showing functional processes enriched in 132 proteins unique to B-ALL samples studied.
Data information: Each circle represents a non-redundant cluster term. Circle size and colour are based on level of significance (-Log$_{10}$ p-value). Clusters comprise connected enriched terms, and the most representative category is highlighted (additional details in Methods).

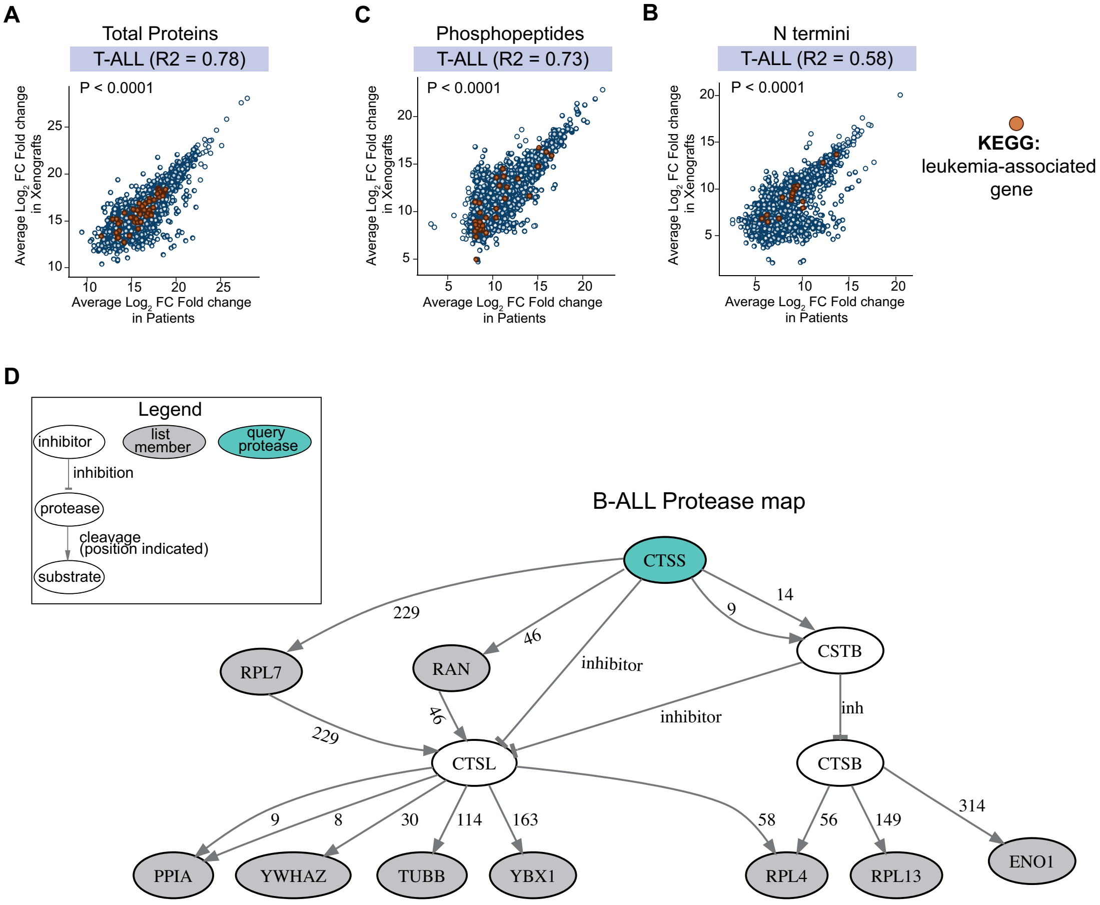

**E**

| Duplicate PDX correlation score | |
|---|---|
| **B-ALL** | **Mean (SD)** |
| B-03-Dx | 0.87 (0.05) |
| B-02-Dx | 0.87 (0.03) |
| B-06-R3 | 0.91 (0.06) |
| B-07-R3 | 0.87 (0.03) |
| **T-ALL** | **Mean (SD)** |
| T-01-R1 | 0.65 (0.02) |

*No data for B-05-R1

**Uzozie et al., Supplementary Figure 9**

**Supplementary Figure 9 - Protein, phosphoprotein and N Termini stability in PDXs**

A, B, C. Correlation of average Log2 fold changes between patients and PDXs for (A) Proteins (N = 5554), (B) Phosphopeptides (N = 2973) and (C) N termini (N = 2832) in all T-ALL patients and xenografts. Proteins annotated to KEGG leukemia-associated genes are coloured in dark brown filled circles for each group of proteins (N = 54), phosphopeptides (N = 35) and N Termini (N = 17).

D. Protease web plot of CTSS interaction network in B-ALL. CTSS (query protease), and list members (N = 235) are indicated. The amino acid following the protease N terminal cleavage site (amino acid P1') is specified for each protease-substrate path.

E. Spearman correlation score comparing OCCRA protein intensities in multiple PDXs expanded from the same patient. Mean score is indicated with the standard deviation from the mean in brackets.

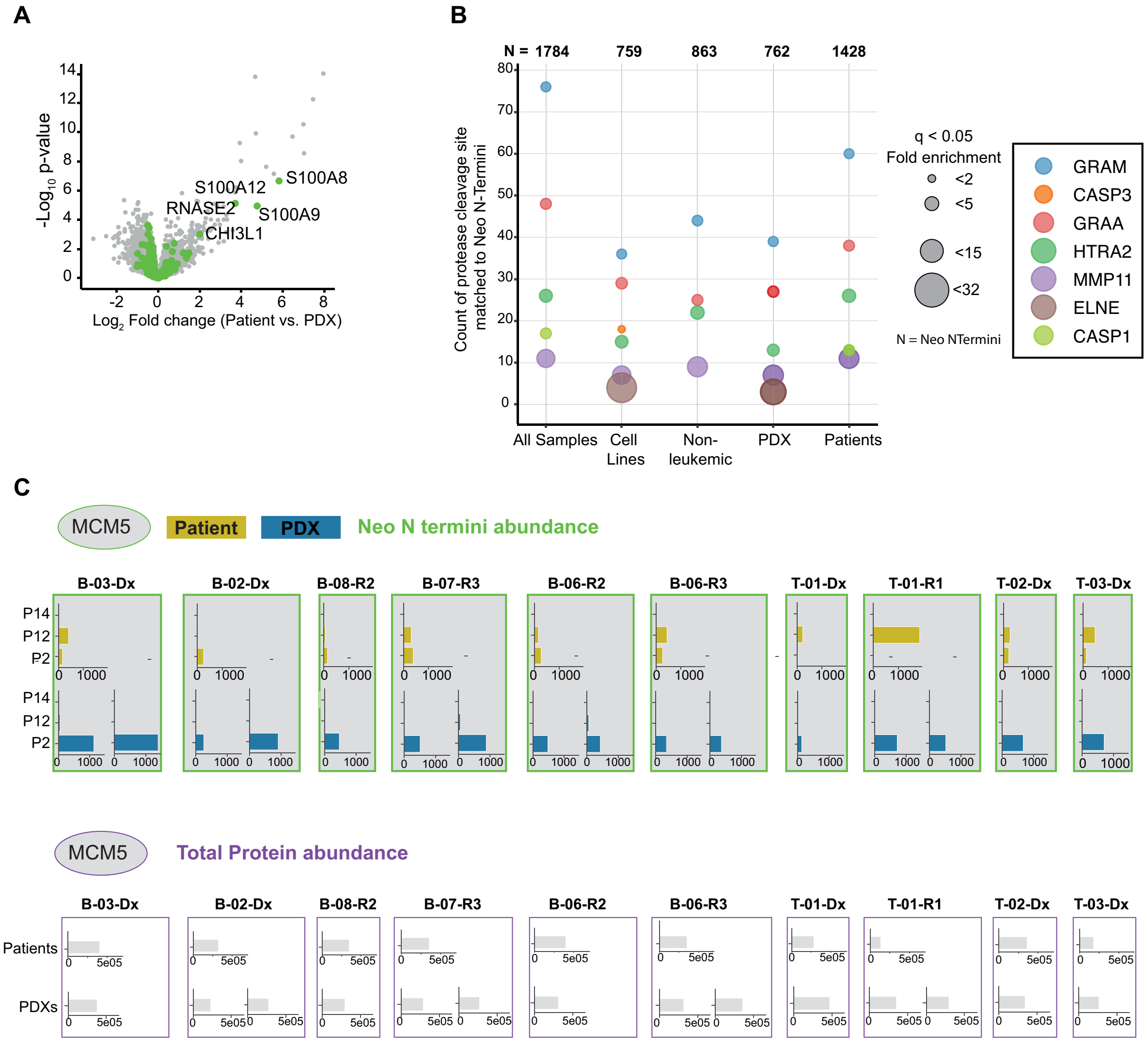

**Uzozie et al., Supplementary Figure 10**

**Supplementary Figure 10 - Similarities and differences in chemokines, proteases, and protein N termini between patients and PDXs.**
A. Comparison of abundance changes in proteins involved in chemokine and cytokine production in patients and PDXs.
B. Annotated significantly enriched proteases (q value < 0.05) and number of protease cleavage patterns enriched in N termini quantified in cell lines, non-leukemic patients, PDXs, and patient ALL samples, in comparison to all samples. Proteases are represented by circles. The size of a circle indicates the fold enrichment factor for a protease.
C. MCM5 proteolytic pattern following GZMM and HTRA2 cleavage in pediatric ALL patients (yellow bars) and corresponding PDXs (blue bars). Average N termini intensities in paired patients and PDXs are shown. P2 (mature protein N termini), P12 (neo N termini) and P14 (neo N termini) indicate protease cut-sites. MCM5 protein intensity in patient and matched PDXs is shown in purple boxes.

### Supplementary Tables

Supplementary Table 1
Supplementary Table 2

### Supplementary Dataset

Supplementary Dataset 1
Supplementary Dataset 2
Supplementary Dataset 3
Supplementary Dataset 4
Supplementary Dataset 5
Supplementary Dataset 6
